# Elongation during segmentation shows axial variability, low mitotic rates, and synchronized cell cycle domains in the crustacean, *Thamnocephalus platyurus*

**DOI:** 10.1101/270728

**Authors:** Savvas J Constantinou, Nicole Duan, Ariel D. Chipman, Lisa M. Nagy, Terri A. Williams

**Author notes:** ***Author Present Address:*** Savvas J. Constantinou- Department of Integrative Biology, Michigan State University, East Lansing, MI, USA, 48824.

## Abstract

Segmentation in arthropods typically occurs by sequential addition of segments from a posterior growth zone, but cell behaviors producing posterior elongation are not well known. Using precisely staged larvae of the crustacean, *Thamnocephalus platyurus*, we systematically examined cell division patterns and morphometric changes associated with posterior elongation during segmentation. We show that cell division is required for normal elongation but that cells in the growth zone need only divide ~1.5 times to meet that requirement; correspondingly, direct measures of cell division in the growth zone are low. Morphometric measurements of the growth zone and of newly formed segments suggest tagma-specific features of segment generation. Using methods for detecting two different phases in the cell cycle, we show distinct domains of synchronized cells in the posterior. Borders of cell cycle domains correlate with domains of segmental gene expression, suggesting an intimate link between segment generation and cell cycle regulation.

**Summary Statement:** Posterior growth zone has synchronized cell cycle domains but shows little cell division during segment addition in a crustacean. Dimensions of the shrinking posterior growth zone change at tagma boundaries.

## Introduction

Arthropods are the most diverse phylum on earth, and much of that diversity derives from the variability in their segmented body plan. This variability includes segment number, size, and character. The developmental mechanisms that produce segments have been extensively studied in the model organism, *Drosophila melanogaster*. But *Drosophila* is atypical among arthropods in that it establishes segments simultaneously, through progressive spatial subdivision of the embryo (Lawrence, 1992). By contrast, the vast majority of arthropod species add their segments sequentially, from a posterior region termed the “growth zone”. In contrast to *Drosophila*, these species elongate as they add segments, thus posing fundamental questions not addressed by the model system: How does elongation occur in the posterior? How and to what degree are elongation and segmentation integrated (Chipman, 2008)? While some mechanisms of elongation are known (*e.g*., teloblastic growth in malacostracan crustaceans; Scholtz, 1993), surprisingly little is known about the range of cell behaviors (*e.g*., cell division or cell movement) responsible for elongation throughout arthropods.

Because most species elongate significantly during segmentation, classical concepts of posterior growth generally invoke mitosis, either in posterior stem cells or in a vaguely defined posterior region of proliferation (Snodgrass 1938; Anderson 1967; Anderson, 1973; Davis and Patel, 2002; Liu and Kaufman, 2005). Cell movement has also been assumed to play a role in elongation in cases where embryonic shape changes dramatically (Tautz et al.,1994; Liu and Kaufman, 2005) - and is documented in some cases (Nakamoto et al., 2015) - but mitosis remains a key driver. Despite these assumptions, cell division and cell movement have rarely been systematically examined: it is simply unknown what drives elongation and exactly how much “growth” is required in the growth zone (Tautz et al.,1994; Davis and Patel, 2002; Liu and Kaufman 2005). This lack of documented posterior growth has prompted some researchers to reject the idea and designate that region the “segment addition region” to avoid confusion (Janssen et al., 2010).

In contrast to our lack of understanding of cellular mechanisms of elongation, the models of the gene regulatory networks that pattern segments in sequentially segmenting arthropods are expanding (reviewed in Williams and Nagy, 2017; Auman and Chipman, 2017; Damen 2007; Peel et al., 2005). It is well established that posterior *Wnt* signaling establishes a posterior gradient of the transcription factor *caudal* (*cad*), which, through downstream genes, progressively subdivides the anterior growth zone and eventually specifies new segments (reviewed in McGregor et al., 2009; Williams and Nagy, 2017). In some systems, posterior *Wnt* signaling is also thought to keep cells in a pluripotent state, presumably dividing as needed to fuel elongation (Chesebro et al., 2013; Shinmyo et al., 2005, McGregor et al., 2009; Auman et al. 2017). The precise links between segmental patterning and cell behaviors responsible for posterior elongation (*e.g*., division, intercalation/movement, shape change) are only inferred, in part, because the cell behaviors themselves remain poorly described for many species. For example, are there many mitotic cells and are they restricted to the growth zone, or do cells from other areas significantly fuel posterior elongation? To fully understand the development and evolution of arthropod segmentation, a more detailed understanding of the cellular mechanisms by which arthropods elongate and grow is needed (Peel et al., 2005). That information will provide vital input to our interpretations of function *via* knock-down/knock-out studies.

Our approach has been to quantify the changes in the growth zone during segmentation in three pancrustaceans as a means of establishing the basis for comparison between taxa: the insects *Tribolium* (Nakamoto et al., 2015), and *Oncopeltus* (Auman et al., 2017); and the crustacean described here, *Thamnocephalus platyurus. Thamnocephalus*, commonly named fairy shrimp, live in temporary freshwater ponds (Rogers, 2009). Their life cycle includes desiccation-resistant encysted eggs, giving rise to commercially available cysts for study (primarily freshwater toxicology, e.g., Alvarenga et al., 2016). After rehydration, cysts hatch as swimming larvae with three pairs of head appendages and an undifferentiated trunk. Sequential segment addition and progressive differentiation gradually produce the adult morphology of eleven limb-bearing thoracic segments and eight abdominal segments, the first two of which are fused to form the genital region (Linder, 1941; Anderson 1967; Fryer 1983; Moller et al., 2004).

In *Thamnocephalus*, we demonstrate that segments are added at a constant, species-specific rate. We characterize the growth zone and newest added segment during segment addition using morphometric measures and find that changes in these measures correlate with position along the body axis, specifically, they occur at tagma boundaries and the position of the first larval molt (*i.e*. between the sixth and seventh thoracic segment). Despite expectations for mitosis to drive elongation in this species, we demonstrate that mitosis in the growth zone is relatively rare; it is required for elongation, just at much lower rates than anticipated. Examination of cells undergoing DNA synthesis (S phase) reveals discrete domains of apparently synchronized cells in the anterior growth zone and newest segments. In *Thamnocephalus*, boundaries of cell cycling domains correlate precisely with *Wnt* and *cad* expression in the growth zone, suggesting direct regulation of these behaviors by the segmentation gene regulatory network.

## Results

### *Segment addition and morphogenesis occur progressively in* Thamnocephalus *larvae*

*Thamnocephalus* hatches with three differentiated larval head appendages (first antennae, second antennae and mandibles) that function in swimming and feeding (Williams, 2007). In addition, the first and second maxillae and approximately three thoracic segments are already specified, as determined by the expression of the segment polarity gene, *engrailed*. As larvae grow, segments are added gradually from the posterior growth zone (Fig. 1) and, likewise, mature gradually. Thus, the trunk typically shows the progression of segment development: segment patterning, segment morphogenesis, and limb morphogenesis (see Constantinou et al., 2016). Segment patterning occurs in the growth zone (defined as the region posterior to the last *engrailed* stripe and anterior to the telson) and is characterized by segmental gene expression but no overt morphogenesis. As is common in sequentially segmenting arthropods, the segments of *Thamnocephalus* form a developmental series along the anterior posterior body axis, with the more mature, anterior segments undergoing limb morphogenesis while new segments continue to be specified in the posterior. Expression of *engrailed* (En) at the anterior of the growth zone indicates that a new segment has been specified. As segments develop, epithelial changes at the intersegmental regions lead to bending of the epithelium and outpocketing of the ventral to ventrolateral surface (Fig. 1C). The initial outpocketing is characterized by a highly aligned row of cells that form its apical ridge. The entire ventrolateral outpocketing eventually forms the limb bud and will begin to develop medial folds along its margin, producing the anlage of the adult limb branches prior to limb outgrowth (Williams, 2007; Constantinou et al., 2016).

**Figure 1.**
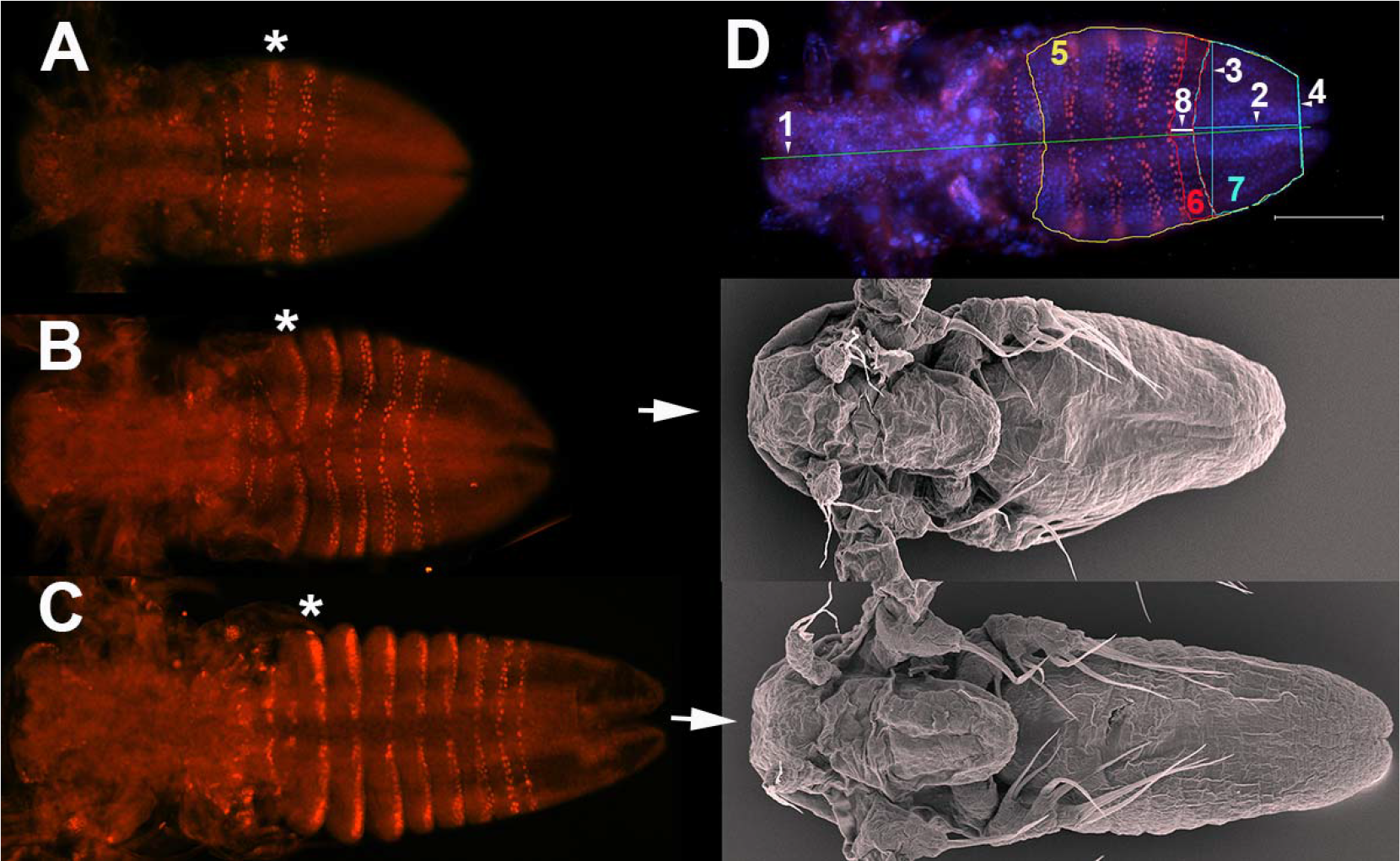
*Thamnocephalus platyurus* development and morphometric measures. A-C. Engrailed protein staining in three different stages of development: A. three En stripes, B. six En stripes, and C. eight En stripes. Note that the asterisks mark the first thoracic segment in each larva and in (C) show the outpocketing of the segmental limb bud form the body wall. In B. and C. scanning electron micrographs of similarly aged larvae show overall morphology. D. *Thamnocephalus* larva with five En stripes, depicting measurements used in this study (defined in methods). Engrailed expression (red) is depicted against cell nuclei (blue-Hoechst). 1- body length, 2- growth zone length, 3- growth zone width “A” (width of newly added Engrailed stripe), 4- growth zone width “B”, 5- trunk area, 6-last segment area, 7- growth zone area, 8- last segment length. Note, the area measures are in color; length measures are given in white and denoted with an arrowhead. Scale bar = 100 μm. All larvae are shown with anterior to the left, ventral side up.

### The rate of segment addition is linear as body length doubles

To characterize the rate of segment addition, we measured the number of segments, as indicated by En stripes, in one hour intervals for staged cohorts of 20-30 larvae. Although we find some variability within each time point, we see a clear trend of linear segment addition (Fig. S1). Segments are added at a rate slightly less than one segment per hour at 30°C. The regularity of segment addition is unaffected by either the first molt (at approximately four hours post-hatching, Fig. S2) or the transitions between addition of thoracic (post-maxillary segments, 1-11), genital (12, 13), and abdominal segments (14-19). During the course of 18 hours at 30°C, *Thamnocephalus* add 14 segments and the overall length of the body roughly doubles (Fig. 2A). Despite the regular periodicity of segment addition, the change in body length at each stage varies, with a noticeable increase following the first molt (Fig. 2B). The overall area of the trunk also increases at successive larval stages (Fig. 2C) and shows a similar per-stage variability.

**Figure 2.**
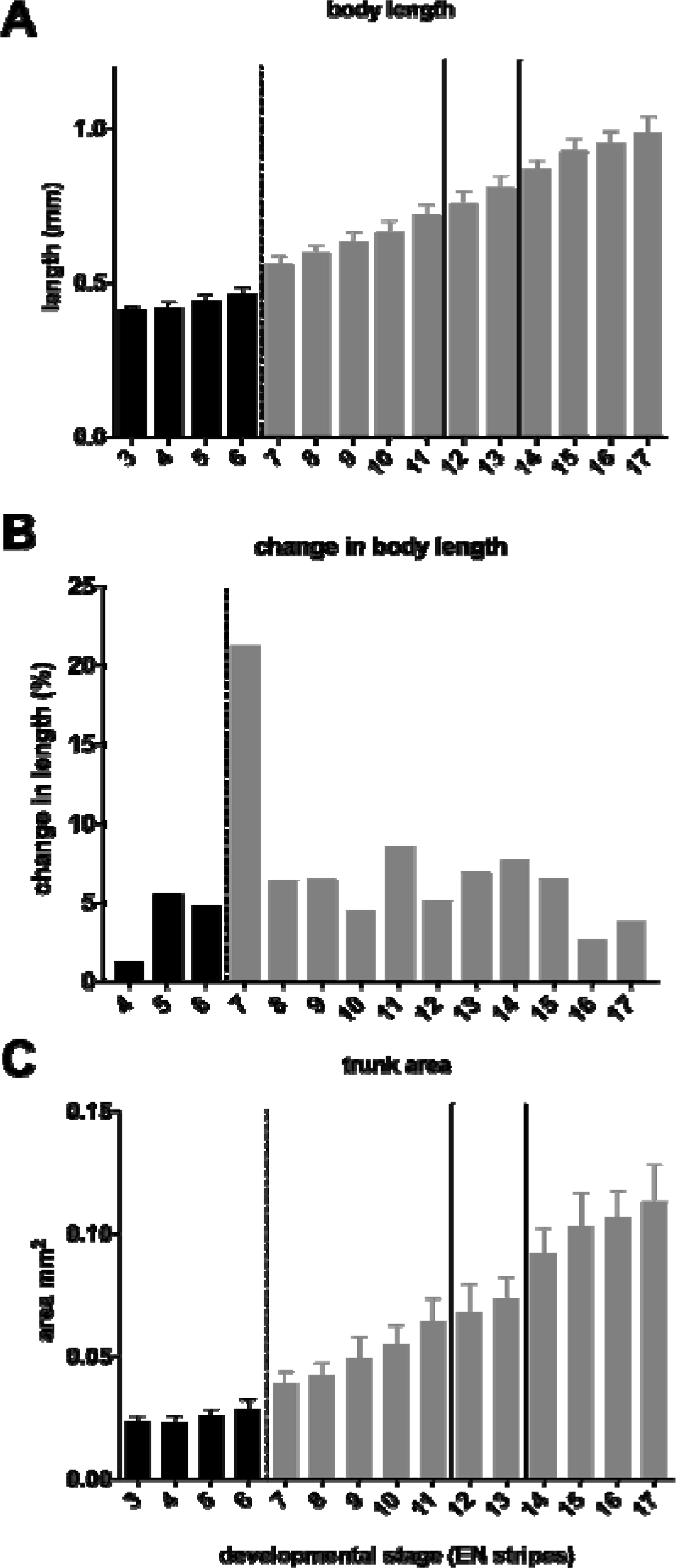
Elongation of the body at successive developmental stages in *Thamnocephalus*. A. Body length plotted against development stage. The animals roughly double in length as the body segments are specified. B. Percent change in body length plotted against developmental stage, demonstrating the impact of the first molt on change in body length. C. Increase in overall area of the trunk during larval development; like the body length, this increases at each stage (after four Engrailed stripes added.) The black bars represent the thoracic segments added before the first molt (dashed line), subsequent thoracic segments are grey. Genital segments (modified abdominal segments 1&2) are marked by solid lines and are followed by additional abdominal segments.

### The growth zone varies during axial elongation and requires growth to produce all segments

To find out if the growth zone itself is changing over time and to understand the requirement for growth as segments are being added, we measured several features in each stage (Fig. 1D; including the length, width, and area of the growth zone and last added segment; as well as, the overall body length and trunk area). In general, there is a decrease in most *Thamnocephalus* growth zone measures as segments are added (Fig. 3). Both the length and the area of the growth zone decrease over time. The exception to this trend occurs at the first molt, between approximately 6 and 7 En stripes or around 3.75 hours (30°C; dotted lines Fig. 3, Fig. S2). Post-molt, the growth zone increases in length (Fig. 3A,B) and area (Fig. 3D), which would be expected after release from the cuticle. Although the overall trend of a successively depleted growth zone might appear intuitively obvious, it is not a given; in the related anostracan branchiopod, *Artemia franciscana*, we found that the growth zone maintains its size through the addition of the first 9 En stripes (Fig. 3C). The general decrease in the *Thamnocephalus* growth zone size plateaus as the thoracic segments are added and then again when abdominal segments are added (see Fig. 3D, E, G; tagmata are separated in the graphs by solid lines).

**Figure 3.**
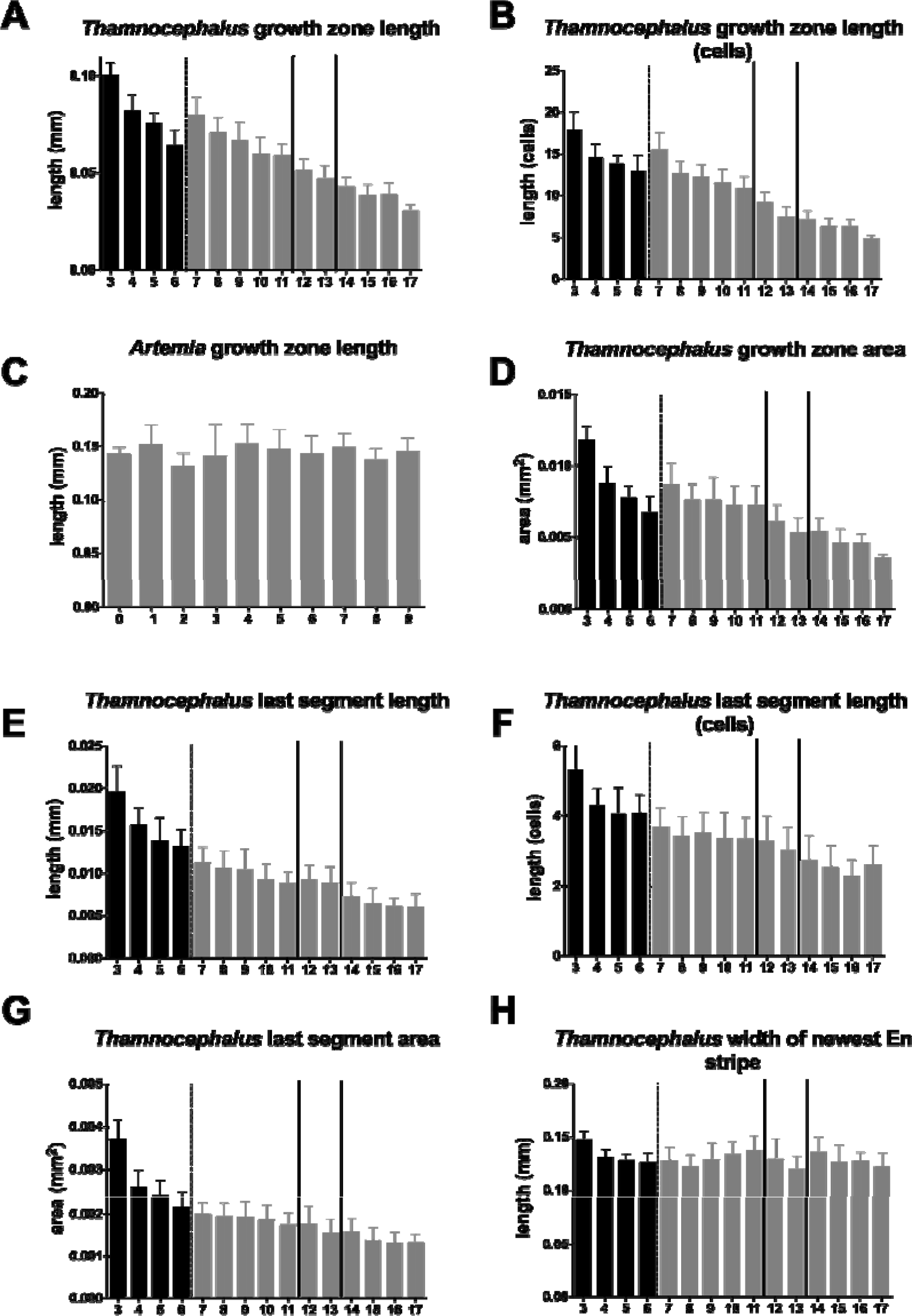
Change in growth zone dimensions in growing *Thamnocephalus* (A, B, D-H) and *Artemia* (C) larvae. A. Length of the growth zone at successive developmental stages in *Thamnocephalus* shows the growth zone shortens over time except when it lengthens after the first molt. This trend is the same when measured by counting cells (B) but indicates how few cells are in the growth zone, especially at later stages. C. In contrast to *Thamnocephalus*, the length of the growth zone at successive developmental stages in *Artemia* shows the growth zone length does not vary much over time. D. In *Thamnocephalus, t*he area of the growth zone decreases, except after the first molt. E. The newest segments are greatest in length during early stages. F. When measured by counting cells, the length of the newest segment added mimics the linear dimension in (E) but shows how few cells make up each new segment anlage, especially at later larval stages. G. The area of the last added segment decreases over development in *Thamnocephalus*. H. Unlike other dimensions, the width of *Thamnocephalus* larvae where the newly specified Engrailed stripe forms remains relatively constant during development (growth zone width “A” measure). The black bars represent the thoracic segments added before the first molt (dashed line), subsequent thoracic segments are grey. Genital segments (modified abdominal segments 1&2) are marked by solid lines and are followed by additional abdominal segments.

To examine the significance of tagmata during segment addition, axial positions were split into four groups for statistical analysis, a measure’s “tagmal designation” was defined by the position along the body axis of the last added *engrailed* stripe: *engrailed* stripes 3-6 = thoracic pre-molt; 7-11= thoracic post-molt; 12-13 = genital; 14-17 = abdominal. We find that axial position is significant in most morphometric measurements, when individuals are grouped by tagmata and compared (Fig. S3). For example, each tagma forms segments from a successively smaller growth zone, whether measured by length (Fig. 3A, B) or area (Fig 3D). The one measure that remained notably steady between tagmata was the ‘growth zone width A’ measure, which is the width of the last En stripe (Fig. 3H). We further test these trends by analyzing morphometric measurements using Principal Component Analysis (PCA) and find significant differences by axial position (Fig. 4). PC1-PC3 explain 93.4% of the variation in the data and are different by ‘tagma’ (Type II MANOVA; F_9,1272_=103.06, p<0.001). PC1 explains 67.5% of the variance and separates by ‘tagma’; a linear model of PC1 by axial position suggests all groups are significantly different (adj R^2^ = 0.814; p<0.001). Intriguingly, the thoracic segments added pre- and post-molt formed groups that are as distinct as the other ‘true’ tagmata.

**Figure 4.**
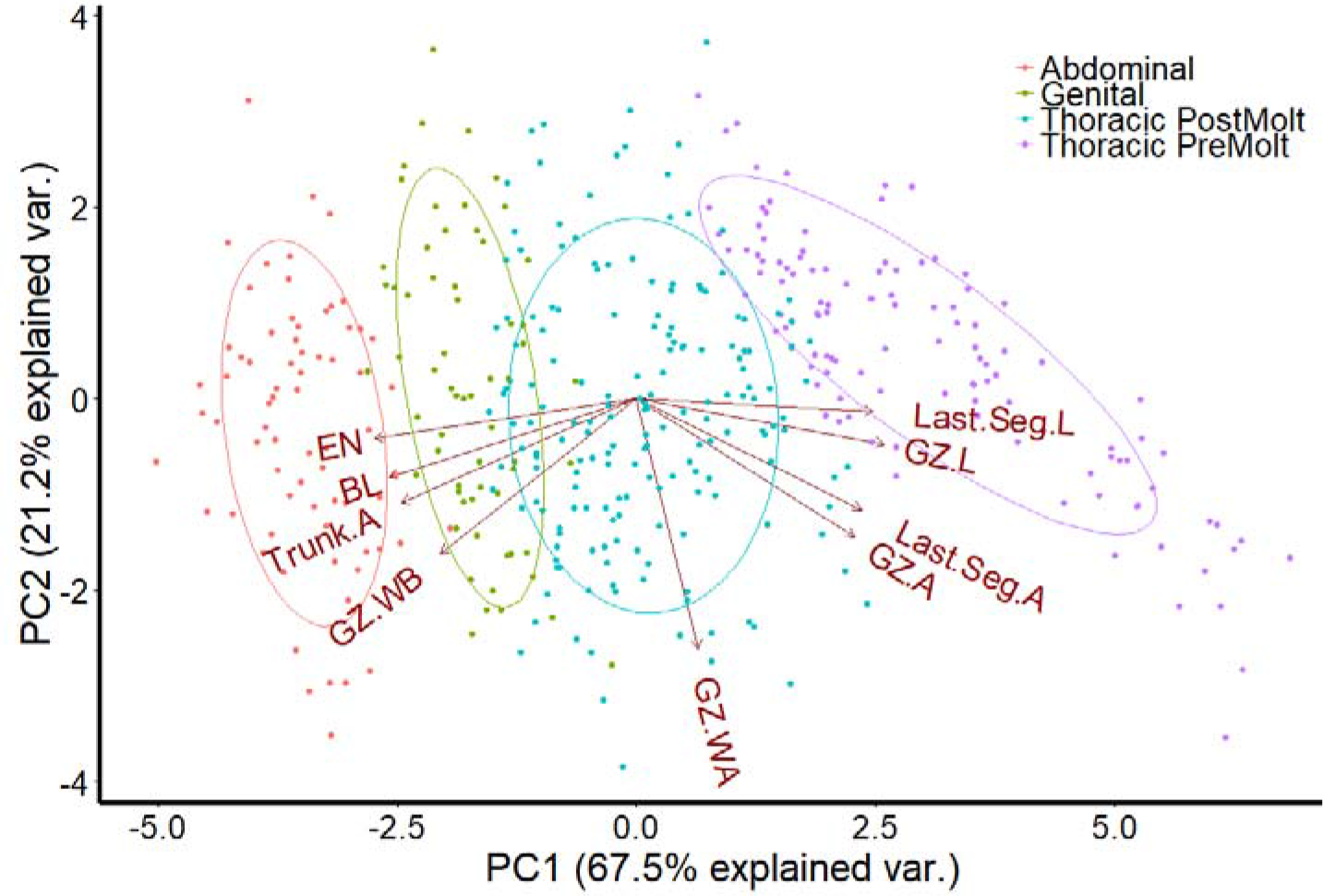
Principal Component Analysis biplot. 423 individuals are plotted along PC1 and PC2 and colored by tagma (the tagma of the last segment formed). PC1 separates individuals by tagma with each tagma group being significantly different from one another. In addition, thoracic pre- and post-molt segments form clusters that are significantly different from all other tagma (Type II MANOVA; F_9,1272_= 103.06, p<0.001).

In addition to linear measures, we counted numbers of cells (nuclei) along our measured linear dimensions, to characterize growth zone changes based on the biological unit of cell number. As can be seen, (compare Fig. 3B to A, and Fig. 3F to E), the trends based on cell counts parallel those of linear dimensions, showing that during this phase of segmentation there is little change in cell shape that would skew the two different types of measurement (linear dimension and cell counts). However, the cell counts are interesting because they describe the changes in terms of the biological unit of cellular dimensions. For example, we find that the smaller segments that are added posteriorly are only 2-3 cells long as compared to 5-6 cells long in the first segments added.

During the time we tracked segment addition, approximately 14 segments were added. The body length increased about 140%, from 0.41 mm to 0.98 mm (Fig. 2A). The total area of the 14 added segments - when measured just as each is being formed in successive stages - represents an area 22 equal to 0.029 mm^2^. The area of the initial (hatchling) growth zone is 0.0118 mm^2^ or only about 40% of the total area ultimately needed to add all the segments (Fig. S4). During segmentation, the growth zone shrinks (Fig. 3A, D) but even a completely depleted growth zone would only account for the addition of approximately the first four added segments. The growth zone needs to more than double to produce the material for new segments; it cannot account for all additional segments without some form of growth.

### The growth zone has few mitotic cells and only a low requirement for growth

The larval epithelium is anchored to its cuticle in *Thamnocephalus*, making significant cell movements unlikely. Thus, to characterize growth in the growth zone, we focused on mitosis. We first measured mitosis in the growth zone by identifying cells clearly in metaphase, anaphase, or telophase using nuclear staining (Hoechst). The highest numbers of mitoses scored in this way were measured immediately following hatching, with an overall trend of fewer mitoses in the growth zone as segment addition continues (Fig. 5A, grey bars). Mitotic numbers increase slightly just prior to and after the first molt (dotted line in Fig. 5A), but overall mitosis counts are low (ranging from about 2 to 13 cells). For Hoechst mitotic counts, we also scored the orientation of the mitotic spindle and found that mitoses in the *Thamnocephalus* growth zone are oriented parallel to the anterior-posterior (AP) body axis. An average of 80% of all cells undergoing division in the growth zone are oriented in the AP direction, regardless of which segment is being produced (Fig. 5B). In some larval stages as many as 90% of mitotic cells in the growth zone are AP oriented. While, mitotic cells in the growth zone are almost always oriented parallel to the AP body axis, mitoses in the anterior, newly specified, segments are generally oriented transversely to the AP body axis (Fig. 5D, not quantified).

**Figure 5.**
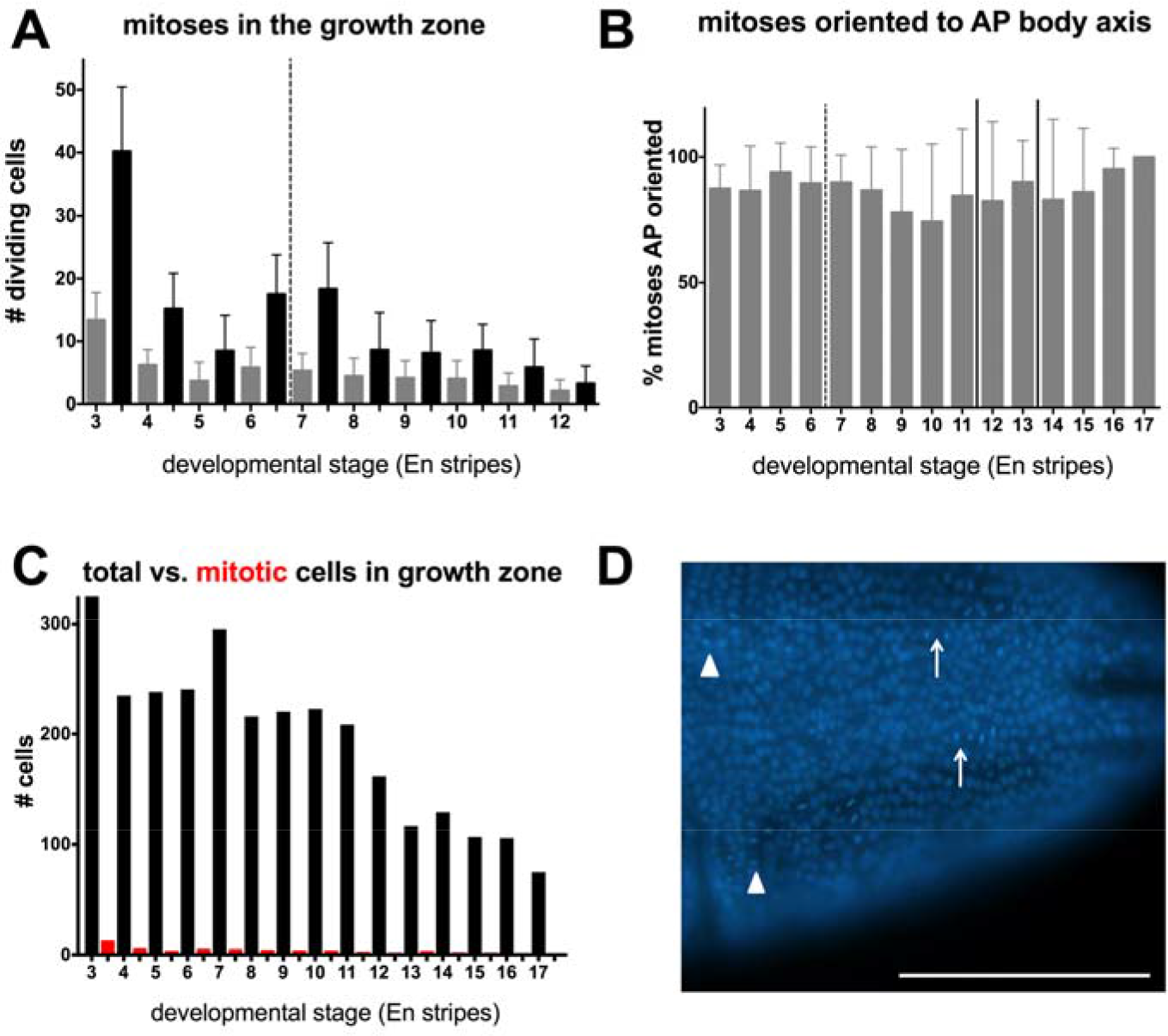
Direct measures of mitosis in the growth zone of *Thamnocephalus*. A. Scoring pH3-positive cells in the growth zone captures consistently higher numbers of cells in M phase compared to cells measured with nuclear staining (Hoechst). Mitosis rates are highest just after hatching and increase prior to the first molt (dotted line). B. Regardless of developmental stage, ~80% of the actively dividing cells in the growth zone, as measured with nuclear staining, are oriented along the AP body axis. C. Total calculated number of cells in the growth zone compared to average number in mitosis at successive developmental stages. D. Representative photo of AP oriented cells in the GZ (arrow). Note the medial-lateral oriented cells in the developing segments (arrowhead). Scale bar equals 100 μm.

To provide a corroborative measure of mitosis, we scored cells that bound to antibodies against phosphorylated histone H3 (pH3; Hendzel et al., 1997). We found that measures of pH3 staining are consistent with the measures obtained by Hoechst and that even the greater number of cells in M phase revealed by this method (Fig. 5A, black bars; 2.4 x more on average). Both measures confirm that total mitotic activity in the growth zone is much less than anticipated by simple overall body elongation. The Hoechst and pH3 measures sometimes showed poor correlation within an individual (Fig. S5). pH3 immunoreactivity is typically initially detected in prophase and fades during late anaphase (Hendzel et al., 1997; Giet and Glover, 2001). While the pH3 signal is required for cells to enter anaphase (Le et al., 2013), the stages of the cell cycle in which pH3 immunoreactivity can be detected vary between species (Hans and Dimitrov, 2001). We found in *Thamnocephalus*, immunoreactivity of pH3 fades prior to anaphase (data not shown). Thus, for any given specimen, cells scored in metaphase, anaphase, or telophase with Hoechst were not always a subset of those scored by pH3 (prophase/metaphase; Supplementary Table 1) and single photographs of either Hoechst or pH3 used to represent typical mitoses may not represent average mitotic rates. Strikingly, even the greater numbers of cells in mitosis revealed by pH3 staining are low relative to the total number of growth zone cells (Fig 5C).

We combined these direct measures of mitosis with our cell counts describing growth zone area to produce estimates of how much division might be required for segment addition. Based on both direct cell counts of growth zone length and width, and calculated cell counts of growth zone area (Fig. S4), the cells in the initial growth zone would need to divide about 1.5 times to produce enough cells to account for the addition of all the new measured segments (14) in this study. While this number seems surprisingly low, it is supported by our direct measures of mitosis compared to total growth zone cell numbers (Fig. 5C): mitotic cells only make up 1-4% of the cells in the growth zone. Consistent with this observation, the area of the larval trunk increases over time (Fig. 2C), much more rapidly than the growth zone or last segment areas decrease, showing that the apparent growth of larvae is disproportionately in the already specified segments, and not in the growth zone *per se*.

### Edu incorporation reveals distinct domains of cell cycling

Mitotic scores in fixed animals give only a snapshot of cell cycle behavior and potentially underestimate rates of cell division. To capture a longer time-course of cell cycling, we exposed animals to 5-ethynyl-2’-deoxyuridine (EdU), a nucleotide analogue incorporated into cells during active DNA synthesis (S phase). A 30 minute exposure to EdU prior to fixation labeled cells actively synthesizing DNA during that time. This method revealed surprisingly stable domains of cell cycling in the larvae (Fig. 6-7).

**Figure 6.**
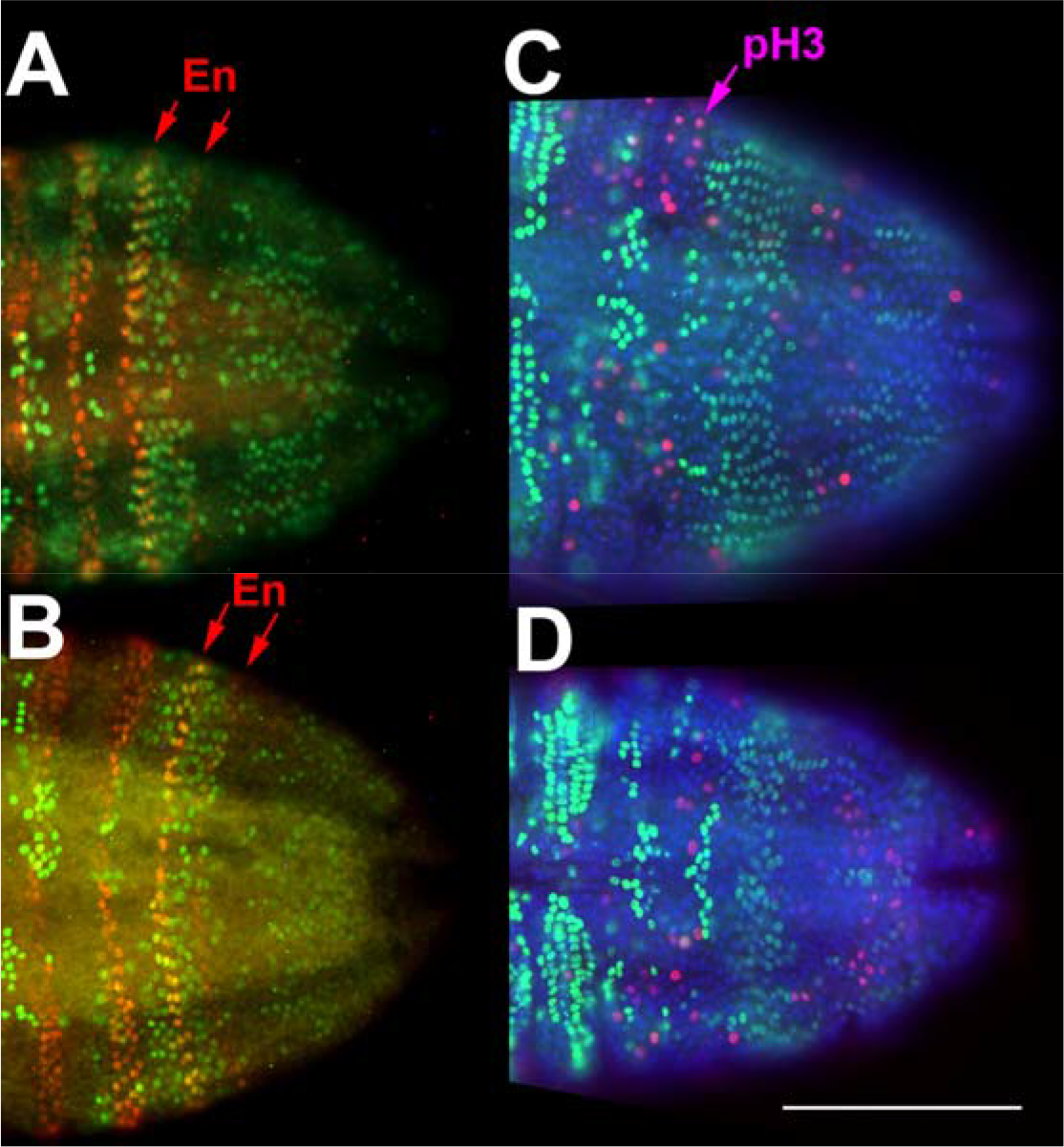
A distinct domain of cells synchronized in S phase appears in the last added segment while the anterior growth zone lacks EdU incorporating cells. A, B. After 30 minutes of exposure to EdU (green), a band of cells in S phase is visible in the last added segment (red arrows indicate last two En stripes) in *Thamnocephalus*. This pattern is maintained throughout the early stages as seen in representative 1 hour (A) and 2 hour (B) larvae. The band lies almost entirely within the last segment after En segment specification. C, D. In both 1 hour (C) and 2 hour (D) larvae, cells in the last added segment (EdU band, light green) do not show pH3 staining (pink) indicative of M-phase. Scale bars equal 100μm.

### The growth zone and newly added segment form three distinct EdU domains

In early larval stages analyzed in detail (0, 1, 2, 3, 4h cohorts), we find a repeated pattern of EdU incorporation that subdivides the growth zone into anterior and posterior domains: the posterior growth zone has apparently randomly positioned cells that are undergoing S phase, while the anterior growth zone is devoid of cells cycling through S phase. Just anterior to the growth zone, in the newest specified segment, all cells undergo S phase synchronously (all cells initiate DNA synthesis within a 30-minute time window). That is, a band of EdU-expressing cells fills the last added segment, someimtes with additional, adjacent cells extending laterally into the penultimate segment (Fig. 6A, B).

Within all staged cohorts, these three domains were present and distinct. The two anterior domains - the EdU synchronous band and the EdU clear band - did not vary. The most posterior domain, where apparently random cells undergo S phase, was more variable. In that region, there are three general classes of EdU incorporation: staining in many growth zone cells (*e.g.*, Fig. 6A), staining in few growth zone cells (*e.g*., Fig. 6D), or bilateral clusters of cells anterior to the telson. Furthermore, in the posterior growth zone, measures of mitosis (pH3) are low compared to the numbers of cells in S phase, suggesting these cells are cycling at low and uncoordinated rates or have variable lengths of time in G_2_. By contrast, cells in the EdU band in the last segment appear synchronous. We tested this by double-labeling specimens with pH3 and EdU to see if there were cells that had entered mitosis within the EdU domain of the last specified segment. We found that pH3-positive cells are virtually always excluded from this EdU domain, suggesting that cells within the domain are truly synchronizing their behavior at the anterior growth zone/newly specified segment boundary, and are maintained in a similar phase of the cell cycle (Fig. 6C-D).

### Segments follow a stereotyped pattern of S phase as they develop

In contrast to the three distinct stable domains of the growth zone region, we see stage-specific patterns of S phase (identified through EdU incorporation) in the more anterior specified segments examined at different stage cohorts. As the animals add new segments from 0 to 4 hours, cells in the anterior segments begin to successively enter S phase (Fig. 7, higher magnification in Fig. 7C, D). This occurs because each segment goes through a stereotyped pattern of S phase cycling as it develops (Fig. 7A, B): first all cells in the segment are in S phase (when the segment is first specified), then no cells are in S phase except those of the neuroectoderm; then S phase is initiated in cells at the apical ridge of the ventral outpocketing segment (in cells that express *Wnt1*, and other *Wnt* genes, just anterior to En; Constantinou et al., 2016); S phase then spreads into other cells throughout the segment.

**Figure 7.**
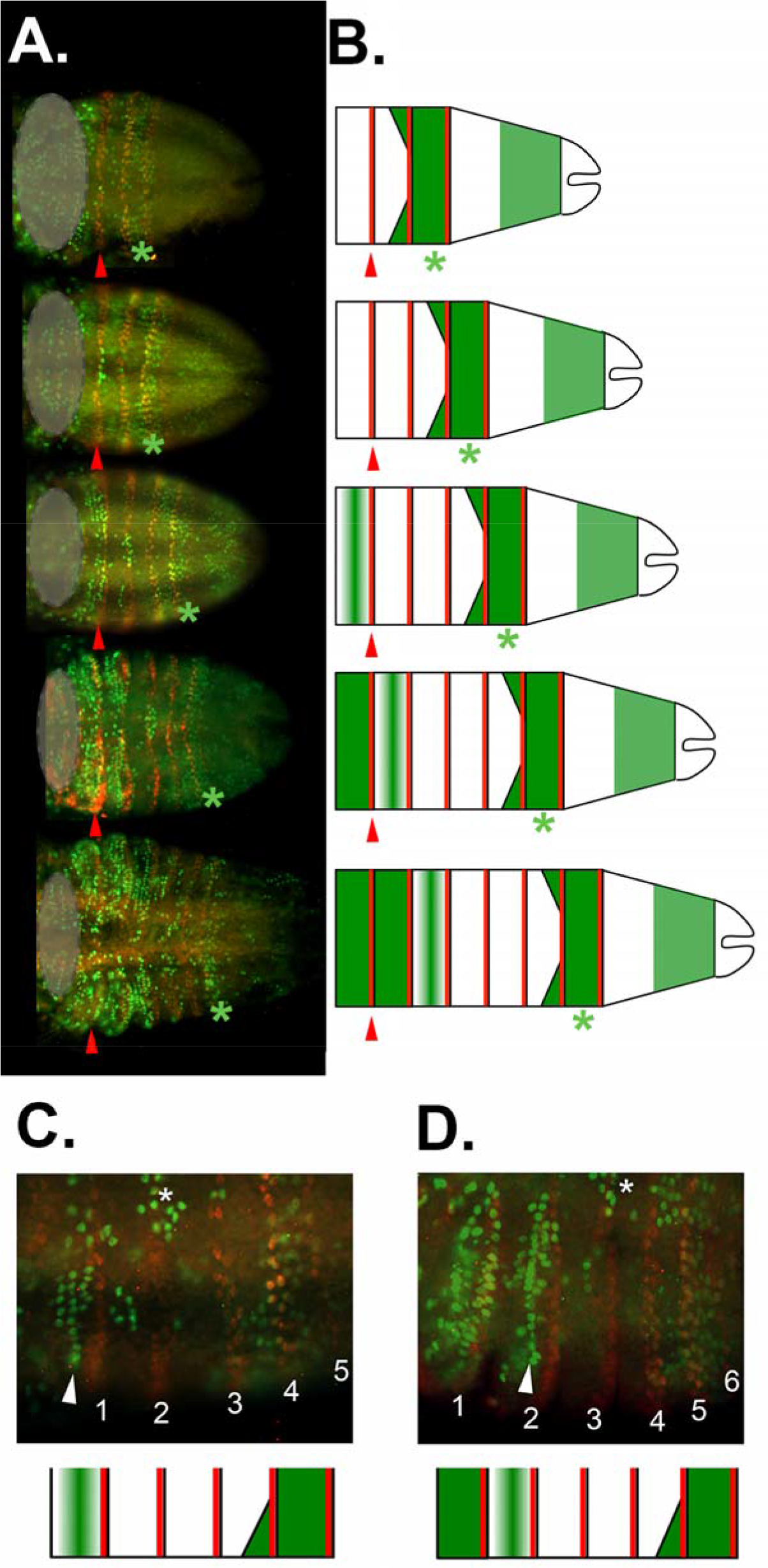
EdU incorporation in anterior segments shows stereotyped progression in *Thamnocephalus* larvae. A. Representative larvae with three to seven segments, oriented anterior left; the trunk is everything posterior (right) of the gray circle (which covers the head segments for clarity). B. Diagrammatic representation of animals with three to seven segments highlighting the progression of EdU incorporation in the trunk of *Thamnocephalus*. A, B. In each stage, the first thoracic segment is denoted by the red arrowhead and the EdU band is indicated with a green asterisk. The anterior growth zone is devoid of EdU, while the posterior growth zone has variable numbers of cells incorporating EdU. In the last added segment, all cells incorporate EdU, forming a band of EdU that sometimes extends into the lateral edges of the penultimate segment. The two segments anterior to this are devoid of EdU. Anterior still, segments begin to progress through S-phase, beginning as a discretely aligned row of cells at the apical ridge of the segment that then expands throughout the segment. C, D. Higher magnification of a series of hemi-segments to illustrate progression of EdU incorporation in the trunk. Thoracic segments are indicated by number and the EdU incorporating cells aligned along the apical ridge are indicated (arrowhead). The neuroectoderm cycles through S phase a few segments anterior to the EdU band (asterisk). Both a specimen (top) and corresponding diagrammatic representation (bottom) are given.

Thus, the overall, appearance at any larval stage depends on the number of segments specified. In 0 hour animals, the two relatively small maxillary segments anterior to the thorax show high levels of EdU incorporation, although thoracic segments one through three, which are already expressing segmentally iterated stripes of En, do not. As animals get progressively older (1-4 hours post hatching) and add more segments, the pattern of anterior segments undergoing S phase continues towards the posterior (Fig. 7).

### Domains of cell cycling in the growth zone correspond to boundaries of segmentation gene expression

We analyzed expression of *caudal* and *Wnt* genes relative to EdU incorporation in the posterior, looking specifically at three *Wnts* shown to have staggered expression in the growth zone: *Wnt6, WntA*, and *Wnt4* (Constantinou et al., 2016). Expression of *cad* is non-graded and extends throughout the growth zone to the border with the telson (Fig. 8A). *WntA* and *Wnt4* expression form two exclusive domains within the growth zone, WntA in the anterior and Wnt4 in the posterior (Constantinou et al., 2016). Strikingly, the domains of Wnt expression map precisely to the domains of EdU incorporation in the growth zone: *WntA* expression in the anterior corresponds to cells lacking EdU incorporation (Fig. 8B) and *Wnt4* in the posterior corresponds to cells with scattered EdU incorporation (Fig. 8C). More anteriorly, the last two stripes of *Wnt4* expression, *i.e*. the most recently formed, appear to flank the band of coordinated EdU positive cells (Fig. 8C). The anterior border of both *cad* and *WntA* also coincides with the posterior border of the EdU domain in the newest segment. Posterior *Wnt6* expression is restricted to the telson, that is, behind the region of relatively dense cells that make up the posterior growth zone (Fig. 8D). Interestingly, cells that form the apical ridge of the limb bud that express *Wnt6* are also those cells that show the early EdU incorporation apically (Fig. 8E).

**Figure 8.**
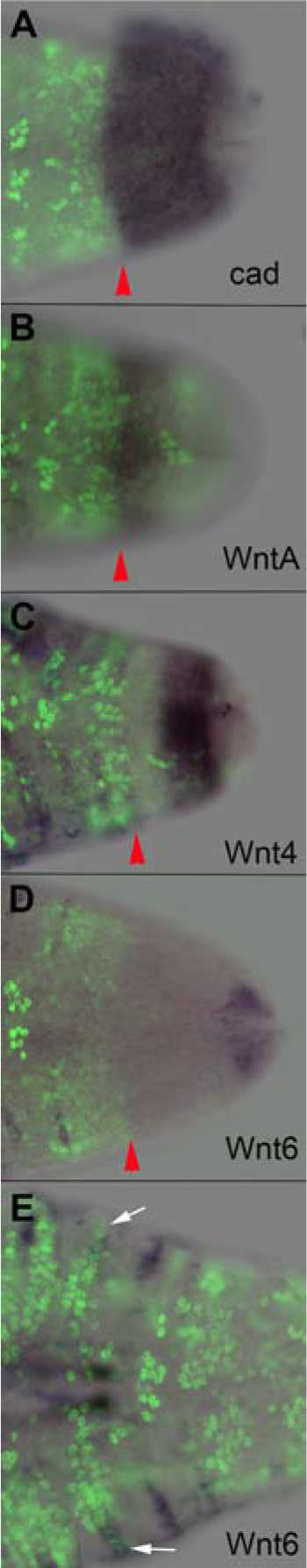
*Caudal* and *Wnt* gene expression maps directly to boundaries of EdU domains. A-E. Posterior of larvae showing both *in situ* expression domains and EdU incorporation. In each case, anterior is left and the posterior edge of the EdU band is denoted by a red arrowhead. A. *Cad* expression extends throughout the entire growth zone and borders the telson, overlapping the posterior *Wnt4* and *WntA* expression. B. Posterior *WntA* expression is mainly in the anterior growth zone, where there are no EdU (or very few) positive cells. The anterior border of *cad* (A) and *WntA* (B) both flank the posterior edge of the synchronized EdU band in the newest specified segment. C. Posterior *Wnt4* expression excludes the band with rare EdU staining and overlaps with the unsynchronized EdU region in the posterior growth zone. *Wnt4* also appears to have a concentration gradient from posterior border towards anterior border. The anterior border of *Wnt4* expression meets the posterior border of *WntA* expression. D. *Wnt6* is expressed in the telson and (E) in the cells that form the apical ridge of the limb buds, which also show EdU expression (white arrows).

## Discussion

### Is there growth in the “growth zone”?

In sequentially segmenting arthropods, axial elongation appears coupled to segmentation in a way that supports the assumption that posterior segmentation is linked to posterior growth. This assumption has been both explicitly recognized (Davis and Patel, 2002; Peel et al., 2005) and challenged (Janssen et al., 2010), leading to a more neutral description of the posterior as a segment addition region and not a growth zone. Furthermore, it is clear in some insects that classical views of a proliferative posterior growth zone are inadequate to explain changes in embryo shape that can accompany segmentation during embryogenesis, and that cell movement plays a significant role in some cases. These cell movements can drive rapid elongation as live imaging and clonal analysis have begun to show, (for example, *Drosophila*, Irvine and Wieschaus, 1994; *Tribolium*, Benton et al., 2013; Nakamoto et al., 2015). Nonetheless, the phenomena responsible for posterior elongation remain understudied, especially compared with the exploration of patterning genes regulating segmentation. They have been studied systematically in two insects - *Tribolium* (Nakamoto et al., 2015) and *Oncopeltus* (Auman et al., 2017) - both of which show a limited requirement for growth. Here, we used careful staging to examine larvae of the crustacean *Thamnocephalus*, whose mode of segmentation appears to have a more obvious requirement for posterior growth than typical insect embryos since segmentation occurs in an epithelium that is anchored to an overlying cuticle, not permitting extreme changes of larval shape. The need for growth could be met by high levels of posterior mitosis, as is assumed for a canonical growth zone (Mayer et al., 2010).

Matching the expectation for growth, we document a ~140% increase in body length during segment addition in *Thamnocephalus*. Unexpectedly, however, systematic examination of mitosis in the growth zone - using conventional means of Hoechst staining or pH3 immunohistochemistry - itself revealed only <5% of cells in mitosis at any given stage. To assess the significance of these mitosis counts strictly with respect to a requirement of posterior growth *versus* overall larval growth, we restrict our calculations of growth to the new tissue that forms each segment as it is added. Based on measures of new segment area and calculations of the area of the initial field of cells that makes up the growth zone, we estimate that cells in the growth zone need only divide between 1 and 2 times (~1.5X) to provide enough tissue to form the new segments measured. This calculation is only a rough estimate, but even so, we find it surprisingly low. We emphasize the misleading nature of overall embryo/larval elongation when analyzing the role of the growth zone in forming new tissue for adding segments. Indeed, in the few species in which mitosis has been examined during sequential segmentation (Freeman 1986; Mayer et al., 2010; Rosenberg et al., 2014; Auman et al., 2017; Cepeda et al., 2017; this study), mitosis in the already specified segments is extensive and no doubt contributes extensively to overall elongation. This leads to a false expectation of high mitosis in the growth zone and at the same time potentially obscures a low but real requirement for posterior growth. Interestingly, our estimates of growth zone cells needing to divide only 1-2 times to meet growth requirements parallels our findings in insects: in *Oncopeltus*, growth zone mitoses were low and their localization revealed only by averaging over a number of staged embryos (Auman et al., 2017); in *Tribolium*, clones of cells labeled in the blastoderm divided about twice prior to germband elongation (Nakamoto et al., 2015; Edgar and O’Farrell, 1990). Our estimate for posterior cell division in *Thamnocephalus* also parallels zebrafish data in which progenitor cells divide only one time after the presomitic mesoderm is established (Bouldin et al., 2014). Average numbers of cells in mitosis in the growth zone do vary by stage, and we find the greatest number of mitotic figures in the growth zone immediately post-hatching as well as just before and after the first molt. The latter is not surprising, since growth *via* molting is often accompanied by mitosis. In summary, despite a measurable requirement for increased area to account for the addition of new segments, the predicted amount of cell division required to make the additional tissue is low and is corroborated by the low counts of mitoses based on direct measures of cells in the growth zone.

### Synchronized cell cycle domains in the anterior growth zone/new segment region map to boundaries of segmental gene expression

The most surprising feature of trying to quantify cell cycling in the growth zone in *Thamnocephalus* arose from exposing larvae to a nucleotide analogue (EdU) to visualize cells in S phase. This unexpectedly revealed distinct S phase domains, demonstrating a kind of spatial coordination in cell cycling not captured by examining mitosis alone. We found two stable cell cycle domains associated with segmentation: a band of cells not undergoing S phase in the anterior growth zone and a synchronized band of cells undergoing S phase in the most recently specified segment. (Cells in the most posterior growth zone are variably in S phase.) The best-known cell cycle domains are the mitotic domains in the embryo of *Drosophila* (Foe, 1989). Among arthropods, we do not know of a comparable case of highly synchronized cell cycle domains in the growth zone *per se*. However, although not apparently as tightly synchronized, we found a similar regionalization of cell division in the growth zone of *Oncopeltus* (Auman et al., 2017): data from staged pH3-stained embryos were combined to generate a heat map of cell division, revealing a region of low cell division in the anterior of the growth zone, and high cell division in the posterior. By contrast, examination of *Tribolium* using EdU exposure showed no apparent regionally distinct incorporation within the growth zone (Cepeda et al., 2017).

To interpret the fixed patterns of S phase domains in *Thamnocephalus*, we followed cell domains mapped to analogous positions in carefully staged larvae. This leads to the following hypothesized sequence of cell behaviors: cells in the very posterior growth zone are undergoing low levels of uncoordinated cycling. Then, as they reach the anterior growth zone, they are coordinated and synchronized, perhaps by a cell cycle arrest. After they are newly specified into a segment, all cells begin to undergo S phase in a synchronized fashion. This entire progression of cell cycling is strikingly similar to that found in zebrafish somitogenesis. In zebrafish, progenitor cells first cycle in the posterior, then arrest in S/G2 as they transit the presomitic mesoderm to form a somite, then begin to cycle again due to upregulation of *cdc25* after somite formation (Boudin et al., 2014). Compartmentalized expression of *cdc25* in the tailbud is required for both extension of the body during somitogenesis and normal differentiation of posterior progenitor cells. We have begun to characterize the cdc25 (*string*) homolog as well as other regulators of cell cycle in *Thamnocephalus* (Duan, Nagy, and Williams, in prep).

We did compare the domains of cells in S phase in *Thamnocephalus* with expression of genes known to regulate posterior segmentation and found that boundaries of gene expression map precisely to boundaries of cell cycling. Both *Wnt* and *cad* are known to function in sequential segmentation in a number of arthropods by maintaining the growth zone and have been hypothesized to maintain cells in a proliferative state (Chesebro et al., 2013; Shinmyo et al., 2005 and McGregor et al., 2009; Hayden et al., 2015). The regulatory interactions in *Thamnocephalus* appear somewhat atypical due to the division of the growth zone by distinct *Wnt* expression, at least compared to the handful of arthropods that have been assayed for multiple Wnts: *Achaearanea tepidariorum* (Janssen et al., 2010), *Strigamia* (Hayden and Arthur, 2014), *Glomeris* (Janssen et al., 2004; 2010), *Tribolium castaneum* (Bolognesi et al., 2008; Janssen et al., 2010), and *Drosophila melanogaster* (reviewed in Murat et al. 2010).

Nonetheless, in all arthropods examined there are distinct regulatory signals in the anterior and posterior growth zone, where commonly *Wnt/cad* signaling in the posterior regulates pair-rule and or *Notch* pathway signaling in the anterior growth zone. Our finding of anterior and posterior regionalization of cell behaviors in the growth zone that map to segmental gene expression is similar to what we found in *Oncopeltus:* the region of low cell division in the anterior of the growth zone is coincident with striped *even-skipped (eve)* and *Delta* expression, *versus* high cell division in the posterior coincident with *cad* and broad *eve* expression. Given the emerging model that posits a regionalization of the growth zone into anterior and posterior domains based on data from the regulatory network specifying segments (Auman et al., 2017), it will be interesting to examine additional species for regionalized cell behaviors in the growth zone that we hypothesize represent a general pattern for arthropod growth zones (Auman et al., 2017).

### *Cell division in the* Thamnocephalus *growth zone is oriented in the anterior/posterior body* axis

In addition to quantifying mitoses in the growth zone, we measured the orientation of mitosis and found that almost all cells are oriented along the AP body axis. AP oriented mitoses can bias growth, impacting elongation *via* cell division, as da Silva and Vincent (2007) demonstrate for *Drosophila* germband elongation. Whether it is important for elongation in other arthropods is unclear. It has been described in *Artemia* (Freeman, 1986), who found as we do AP orientation in posterior cells but oblique and transverse within segmented regions. It has also been described in malacostracan crustaceans, where two rounds of AP oriented cell division in cells budded from the posterior teloblasts establish four rows of cells that form the initial segmental anlage (Dohle et al., 2004; Scholtz, 1996). Given the low rates of mitosis used by *Thamnocephalus*, it is unclear what impact oriented mitosis might have on elongation, *per se*. There could be other functions for oriented cell division, *e.g*. the efficient addition of new segments could be improved by orderly cell arrays, or precise molecular gradients may require cells in a particular orientation. Disrupting regulators of planar cell polarity in the growth zone epithelium could shed light on these potential functions.

### Changes in the growth zone are linked to different body tagma

We document that the growth zone shrinks over time in *Thamnocephalus*: the posterior field of cells is depleted as segments are added. However, this decrease is not simply monotonic, but varies by the particular tagma in which segments are being added. Analysis of various morphometric measures shows that the dimensions of the growth zone as well as the newest segmental anlage are statistically smaller when generating abdominal *versus* thoracic segments. This correlation is intriguing. It is known in vertebrates that extension of the embryo, while a continuous process, relies on different cell populations when forming the trunk *versus* tail (Wilson et al., 2009). The switch from trunk to tail is specifically regulated and mutants in *growth/differentiation factor 11 (Gdf11)* can lengthen the trunk by extending onset of the switch (Jurberg et al., 2013; McPherron et al., 1999). While arthropod segmentation is phenomenologically quite different from vertebrates - relying on the subdivision of an epithelial sheet *versus* specification of motile, mesenchymal cells - we find it intriguing that our measures of the growth zone correlate with tagma boundaries. This suggests that, in arthropods, very early segmental anlage are integrating different patterning signals along the body axis, and may similarly show some switch in cellular behaviors involved with early segment formation in different tagma. In this study we focused on only thoracic segments in more cellular detail but in the beetle *Tribolium*, abdominal segments are formed from much more elongated cell clones than thoracic segments (Nakamoto et al., 2015).

Surprisingly, we also find that thoracic segments formed either before or after the first molt are as significantly different from one another as are between-tagma differences. The cause of this correlation is not entirely clear, there could be a change in mechanical constraint imposed by the cuticle, given that one notable change at the first molt is that the telson elongates; or there could be differences within thoracic segments due to hormonal changes that accompany the molt. To ensure that the hatchling animals as a group (with three thoracic segments and noticeably larger growth zone dimensions, Fig 3.) were not forcing differences in the pre- and post-molt thoracic groups, we conducted an exploratory data analysis of PCA, removing the hatchling individuals. Pre- and post-thoracic groups still separated and were significantly different (data not shown, PCA analysis available on Github).

The morphometric correlations with tagma do not have a corresponding temporal variation: the rate of segment addition is constant in *Thamnocephalus* - under various culture conditions and including nearly all of segment addition (17 of 19 segments), encompassing multiple tagmata. A constant rate is consistent with early measures done in less controlled conditions (Williams et al., 2012) and in the one other branchiopod crustacean in which it has been measured, *Artemia* (Weisz, 1946; Williams et al., 2012). By contrast, in insects, while rarely measured, segment addition rate shows some variability, specifically with tagmata. In *Tribolium*, segmentation rate varies depending on the segments being added: the change occurs at the boundary between thorax and abdomen and correlates with a change in cell movement (Nakamoto et al., 2015). We hypothesized that the slowing of segment addition prior to the rapid addition of abdominal segments was necessary for the extreme cell movements that accompany abdominal segmentation. *Oncopeltus fasciatus*, an insect that only adds abdominal segments sequentially, has a constant segmentation rate and no gross changes in cell movement or behaviors (Auman et al, 2017). Sampling additional insects, where both thoracic and abdominal segments are added sequentially, would increase our understanding of these phenomena, particularly how segmentation rate may change at axial position boundaries. Again, we find changes in segments associated with tagma boundaries at their very inception.

Interestingly, one dimension that remains relatively constant throughout the window we measured is the width of the newly added segment. If the growth zone is roughly a trapezoid with three sides formed by the lateral body margins and the anterior border of the telson, the final side is the width of the last En stripe. Although the overall trapezoidal shape can vary, this is because the anterior telson width and the length of the lateral margins vary, the width of the last En stripe remains the same. Intriguingly this measure was also relatively constant during the sequential segmenting phase in *Oncopeltus* (Auman et al., 2017).

Finally, we note that the absolute dimensions we measured raise some interesting questions about patterning. Over the course of segment addition that we documented, the length of the growth zone shrinks from about 18 to 5 cells long and the newly specified segments decrease in length (5 to 2 cells). If, as we infer from gene expression data, there is a segmentation clock with varying *Notch* and/or pair-rule signaling (personal observation), the oscillations of gene expression are occurring over very different cellular dimensions at different larval stages. It will be interesting to explore the spatial and temporal dynamics of the gene regulatory network that periodically produces new segments as the growth zone shrinks by two thirds.

### Cell cycle domains in anterior segments

Examining EdU incorporation throughout the body in any arbitrary specimen shows only what is apparently a large number of cycling cells. That is, at first glance these patterns of EdU incorporation appear somewhat random and widespread, but strikingly regular patterns of incorporation emerge from comparisons of a number of precisely staged larvae (Fig. 7). During early development, we see a progression of cells undergoing S phase from anterior to posterior in newly specified segments. Over time any one segment goes from no cells in S phase, to neuroectoderm in S phase, to apically located cells in S phase, to most cells in S phase. This suggests a regular progression of cell cycling coupled to the visibly regular progression of morphogenesis in the specified segments (Williams, 2007; Constantinou et al., 2016). Indeed, the synchronization of cell cycle in the early segmental anlage may be needed to accomodate or even drive the subsequent morphogenesis of the limb bud. Freeman et al. (1992) argue that increased cell density (resulting from mitosis) was required for epithelial bending that generates the outpocketed limb bud in the related crustacean, *Artemia*.

Intriguingly, the pattern of Edu incorporation we describe in *Thamnocephalus* pattern bears a striking resemblance to the domains of pH3 expressing cells in the wasp *Nasonia*, that similarly appear to progress from anterior to posterior during embryonic segmentation of successively older embryos (Rosenberg et al., 2014). Rosenberg et al., (2014) document a series of mitotic domains lying exclusively between segmental *eve* stripes (at least in early embryonic stages). Interestingly, Foe (1989) found that the boundaries of mitotic domains in *Drosophila* also corresponded to segmental boundaries (En stripes). Thus, the cell cycle domains in these three species all are tied to segmental boundaries. This kind of domain-specific, timed cell cycling, bespeaks a tightly controlled integration of cell division and segment patterning. The presence of this phenomenon in distantly related arthropods begs for comparative analysis among other arthropod groups to determine if this cell behavior is an ancestral or derived trait.

### A summary of growth zone cell dynamics in *Thamnocephalus*

In *Thamnocephalus*, we find a growth zone that is depleted over time (shrinking cell field) while being replenished by cell division. The amount of cell division in the growth zone is low and the rate of cell cycling appears to be slower in the growth zone than in the newly specified segments. Cell division within the growth zone is aligned along the AP body axis although the impact of this on elongation of the body is predicted to be small relative to the increase in length caused by the rapid growth of segments once they are specified. The growth zone has two distinct domains (Fig. 9): a posterior *Wnt4* expressing region that has some cells undergoing S phase and M-phase and an anterior *WntA* expressing region that has no cells in S phase. Once a segment is specified, the cells of that segment enter S phase in a synchronous fashion. Newly specified segments then undergo a patterned sequence of entering S phase, starting with neuro-ectoderm, then the segmental apical ridge, before spreading broadly throughout the segment, forming an AP pattern of cell cycling along the body axis. While these growth zone features are stable in different stages, other growth zone features change in association with tagma in which segments are produced (*e.g*., linear dimensions). These kinds of cellular dynamics are only beginning to be measured in other species and yet already show a number of intriguing characteristics that may be more widespread among sequentially segmenting arthropods and are likely a source of evolutionary variability underlying the segmentation process: the surprisingly low rates of posterior mitosis, the apparently tight regulation of cell cycle at the growth zone/ new segment border, resulting in coordination or synchronization of cell cycling, and a correlation between changes in the growth zone and tagma boundaries.

**Figure 9.**
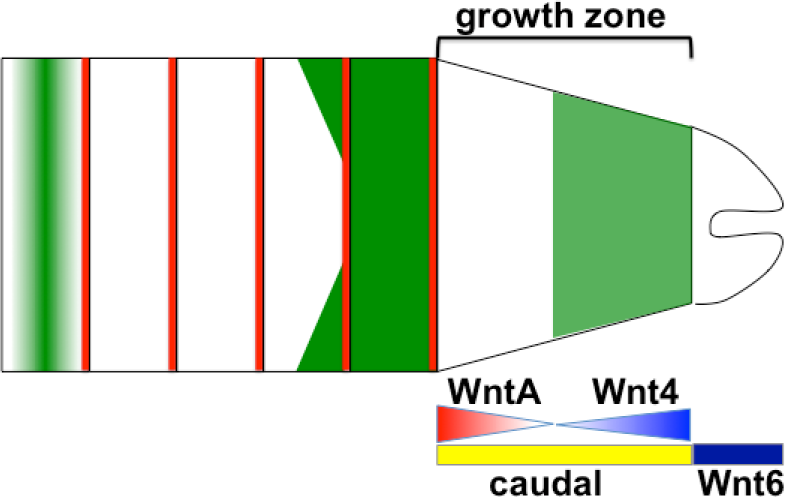
Diagram of growth zone in *Thamnocephalus*. The *Thamnocephalus* growth zone is divided into anterior and posterior regions based on cell behaviors and gene expression. The posterior domain corresponds to *Wnt4* expression (indicated by blue gradient); cell cycling in this region is present but low. Although mitosis in the posterior growth zone is not temporally or spatially synchronized, all mitosis in this domain is restricted in anterior-posterior orientation. The anterior growth zone corresponds to *WntA* expression (indicated by red gradient) and lacks cells in S phase. Cells in this region are possibly arrested either in early S phase or at the entry from G1 to S phase, since immediately after the anterior growth zone cells enter S phase again in the newest specified segment (dark green in last added segment). The synchronized S phase and subsequent mitoses in the segments generate the bulk of the visible elongation of the larvae. *Wnt6* expression (dark blue bar) is in the telson, posterior to the growth zone while *caudal* expression (yellow bar) is throughout the growth zone. S phase domains delineated in green, *Engrailed* expressing cells delineated in red.

## Methods

### Thamnocephalus platyurus *culture and fixation*

*Thamnocephalus platyurus* cysts (MicroBioTests Inc, Belgium) were hatched in 1:8 EPA medium:distilled water solution (EPA medium- 0.0537 mM KCl, 1.148 mM NaHCO_3_, 0.503 mM MgSO_4_, and 0.441 mM CaSO_4_) at pH 7.0 and ~27°C under a full spectrum aquarium lamp (T8 Ultrasun, ZooMed). For precisely staged animals, hatchlings were collected every 15 minutes, raised at 30°C under fluorescent light (~3500 lux) in a Precision 818 incubator. Animals were reared in 6-well cell culture dishes (~5 mL fluid per well) and fed 1 ul of food at time of collection. 4-18H animals received an additional 1 ul of food after a 60% water change at the midpoint of their rearing time while 0-3 hour animals were not fed since they are utilizing yolk reserves. Food consisted of a solution of yeast and commercially available fry food (Hikari First Bites) made fresh each day in 1:8 EPA medium. Animals were fixed for 30 minutes on ice in 9% formaldehyde/ fix buffer (phosphate buffered saline supplemented with 70 mM EGTA) and then dehydrated to 100% methanol in a series of washes (2-3 minutes at 25%, 50%, and 75% methanol). Fixed larvae were stored at 0°C in 100% methanol.

### Artemia franciscana *culture and fixation*

*Artemia* were raised in a 2.5 gallon tank at 25°C, 30-35ppt salinity using artificial sea salts, with continuous aeration and continuous full spectrum light. Newly hatched larvae were collected in timed intervals and were fed a mixture of yeast and algae (see above). Animals were fixed as *Thamnocephalus* (above) but with the addition of 0.1% Triton to the buffer.

### Immunohistochemistry

Immunohistochemistry protocols follow Williams et al. 2002. We visualized Engrailed using En4F11 (gift from N. Patel) and dividing cells using pH3 (anti-phospho-Histone H3 (Ser10) Antibody; Millipore) at 1 μg/mL. Specimens were counterstained with Hoechst, mounted in mounting medium (80% glycerol supplemented with 0.2M TRIS buffer and 0.024M *n*-propyl gallate) using clay feet on coverslips to prevent distortion, and photographed on a Nikon E600 Ellipse epifluorescence microscope and a Spot Insight QE digital camera (Diagnostic Instruments, Sterling Heights, MI, USA) and Spot Advanced software.

### EdU exposures and antibody or in situ doubles

Animals were exposed to 0.6 mM EdU for either 15 or 30 minutes just prior to fixation. EdU was visualized through the Click-iT^®^ EdU Alexa Fluor^®^ 488 Imaging Kit (Thermo Fisher Scientific) as described in the manufacturer’s manual with a final concentration of 1 μM sodium azide. For pH3 doubles, pH3 was then visualized as above. Specimens were counterstained with Hoechst and mounted in 70% glycerol. Photographs were taken as described above. For *in situ*/EdU doubles, animals that had been exposed to EdU 30 minutes prior to fixation first underwent *in situ* hybridization for *caudal* and *Wnt4, WntA, Wnt6* as described previously (Constantinou et al., 2016). Then, after washing out the NBT/BCIP developing solution, animals were washed in 0.1% PBTriton, and processed through the Click-It reaction, as above.

### Molting

Individual animals were collected at hatching (t=0) and allowed to swim freely in 1 mL of pond water in a 24-well plate (Falcon). The timing of the first molt was determined by observing single specimens under a dissecting scope every 5 minutes. The molt was visible as a transparent membrane. Immediately following the molt, the animals also displayed a characteristic behavior: individuals stayed at the bottom of the well and combed the setae on the antennal exopod by repeatedly pulling them between the mandible and gnathobasic spine. After the first molt, the posterior trunk of the animal was elongated compared to the bean shaped trunk before the first molt (Fig. 1 provisional citation) which is reported for other branchiopods (Dahms et al., 2006). The setae on the gnathobasic spine become branched, resembling a bottle-brush, compared to the non-setulated setae before the first molt (Fig. S2).

### Measured and calculated growth zone dimensions

All measurements were made directly on the photographs within the Spot software except number of mitotic cells in the growth zone which were counted in preparations under the microscope. Growth zones measures were confined to the ventral surface since that is the where the active segmental patterning is focused and the growth zone region does not differ materially between dorsal and ventral (Fig. S6). Measures were defined as follows (Fig. 2)

- Body Length (BL): Measurement from the most anterior head region to anus through the midline.
- Engrailed Stripes (En): The number of En stripes posterior to the maxillary stripes. To be scored, the En stripe must extend from the lateral edge of the animal and connect across the ventral surface forming a complete line (i.e., the presence of few, scattered En-expressing cells was not scored as a new segment).
- Growth Zone Length (GZ Length/cells): The growth zone length is measured at the midline from just posterior to the last En stripe to the anterior edge of the telson (which is marked by change in cell density easily seen with Hoechst staining). Cell counts (numbers of nuclei) along this line were also recorded.
- Growth Zone Width “A” (GZ Width A/cells): This measure is from one lateral edge to another just posterior of the final En stripe. The number of cells that make up this measure was also recorded. We refer to this measure as the length of the newly formed Engrailed stripe.
- Growth Zone Width “B” (GZ Width B/cells): This measure extends from the one lateral edge of the posterior growth zone to the other, along the boundary of the growth zone and telson. The number of cells that make up this measure was also recorded.
- Growth Zone Area (GZ Area): This is a roughly trapezoidal measure formed by the two lateral margins of the growth zone and growth zone widths A&B.
- Length Between Two Final En Stripes (Last Seg Length/cells): This is a measurement along the midline of the distance between but not including the final two En stripes. The number of cells that make up this measure was also recorded.
- Last Segment Area (Last Seg Area): This is a measure of the total area of the last segment formed at any specific stage. It is a roughly rectangular measure bounded by the two lateral margins of the segment, growth zone width A and a line just posterior to the penultimate En stripe.
- Trunk Area: This is a measure of the total area of the larval trunk. The measurement includes the lateral edges of all segments and follows the growth zone width B measurement at the posterior. The final portion of the measure is along the second maxillary En stripe, but not inclusive of that stripe. It measures just posterior to the second maxillary En stripe but include the entire area of the first segment.
- Number of Mitotic Cells in Growth Zone: This is a measurement of the number of cells in the ventral epidermis posterior to the last Engrailed stripe undergoing mitosis as visualized by Hoechst 33342 (Thermofisher) or pH3 staining.
- Length and width measures made by cell counts were used to calculate an estimate for the area of the growth zone in cell numbers (using the formula GZ length x ((GZ width A+GZ width B)/2)) as well as cell field area of the last added segment (last segment length x GZ width A). These were used to estimate the number of cell divisions necessary to add all new segments from the initial GZ cell field.

### Statistics

All scatter plots with lines represent linear regressions of the data; all multiple comparisons are done by analysis of variance and show averages with standard deviation. Statistical analyses were performed using GraphPad Prism 7 software or custom R (3.4.0) code. Principal component analysis was conducted with a custom script in R using the ‘prcomp’ function and visualized using the ‘ggbiplot’ package (Vu, 2011). PCA utilized 9 different morphometric measurements (all measures excluding cell counts as outlined in *Growth Zone Dimensions* but also excluding number of mitotic cells) from 423 individuals that were standardized and compared by axial position (tagma). Axial positions were split into four groups for statistical analysis, an individual “tagma designation” was defined by the position along the body axis of the last added Engrailed stripe: Engrailed stripes 3-6 = thoracic pre-molt; 7-11= thoracic post-molt; 12-13 = genital; 14-17 = abdominal.

The following R packages were utilized during data analysis, exploratory data analysis, and visualization; ‘graphics’, ‘devtools’ (Wickham and Chang, 2016), ‘gridExtra’ (Auguie, 2016), ‘data.table’ (Dowle and Srinivasan, 2017), ‘Hmisc’ (Harrell, 2016), ‘extrafont’ (Chang, 2014), ‘broom’ (Robinson, 2017), ‘ggplot2’ (Wickham, 2009), ‘ggsignif’ (Ahlmann-Eltze, 2017), and ‘cowplot’ (Wilke, 2017). All custom R codes and data are available at https://github.com/savvasjconstantinou/tRinityanalysis.

## Acknowledgments

Supported by NSF-IOS 1024220 and 1322350 to T. Williams; NSF-IOS 1024446 and 1322298 to L. Nagy; BSF 2012763 to A. Chipman. Thanks to M. Shankland in whose lab T. Williams first visualized S-phase in fairy shrimp many years ago. Thanks to Dr. Christie Bahlai, Dr. William Pitchers, and Colin Diesh for help with R code, data manipulation, and statistics.

## Competing interests

The authors declare no competing or financial interests.

## Supplementary data file

**Supplementary Figure S1.**
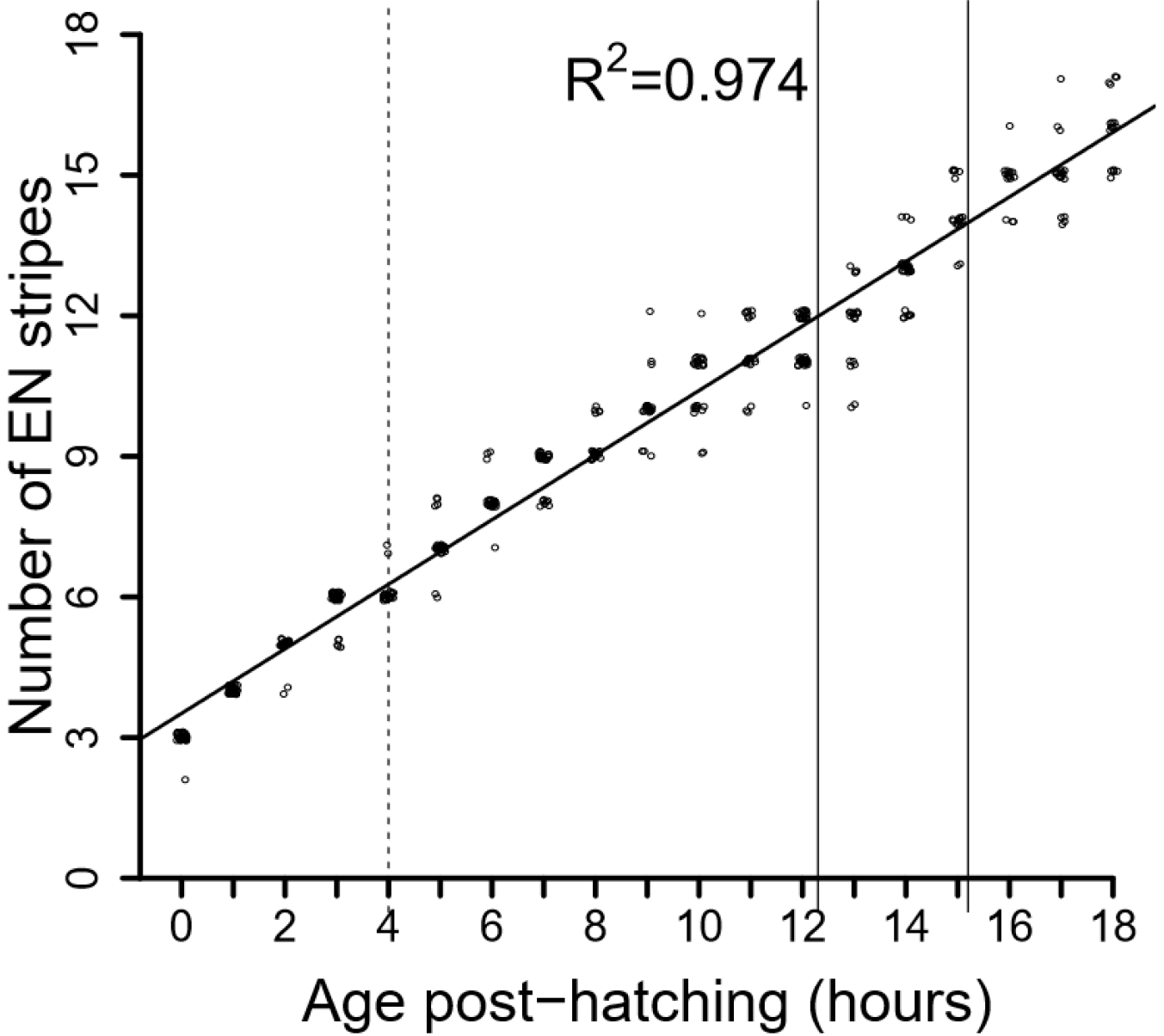
*Thamnocephalus* adds segments linearly. Segment number is plotted against time at one hour intervals and fit with a linear regression. Points are offset to demonstrate the high number of similar measures; n=20-30 individuals for each time point. Dotted line represents the first molting event at 4 hours. Solid lines represent the transition between tagma, thoracic to genital (~12H) and genital to abdominal (~15H).

**Supplementary Figure S2.**
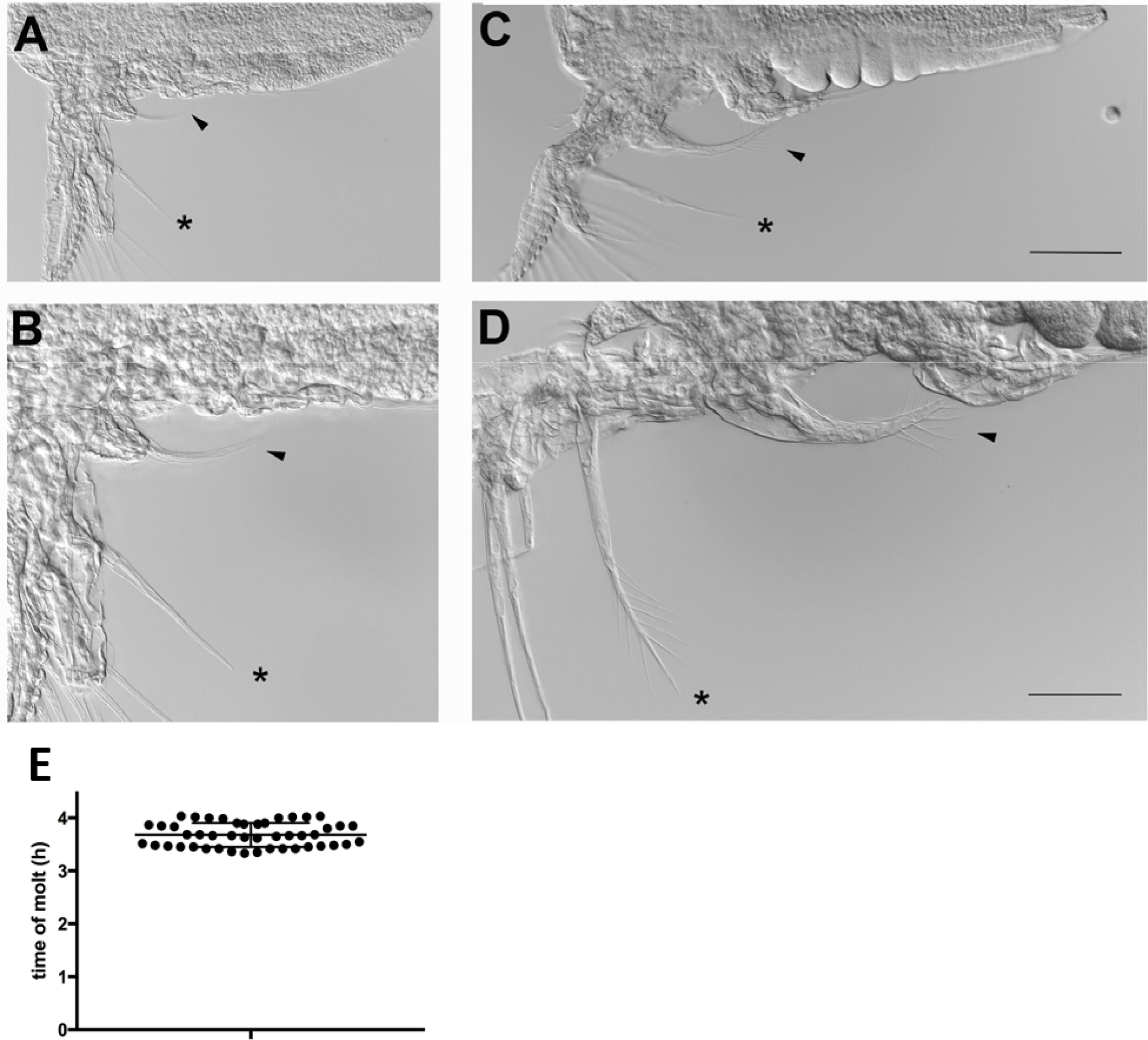
Change in setal morphology that occurs during first molt; used to score animals pre- and post-molt when not tracked as individuals. A, B. Premolt larva showing the relatively smooth trunk (symbol) and the non-setulated gnathobasic process (arrowhead) and xxx spine (asterisk). C, D. Post-molt larva showing overt trunk morphogenesis in the anterior segments (symbol) and the setulation of the gnathobasic process (arrowhead) and xxx spine (asterisk). Scale bars = 100 um. E. Average (3.7h) and standard deviation of time to first molt for a cohort of 46 hatchlings.

**Supplementary Figure S3.**
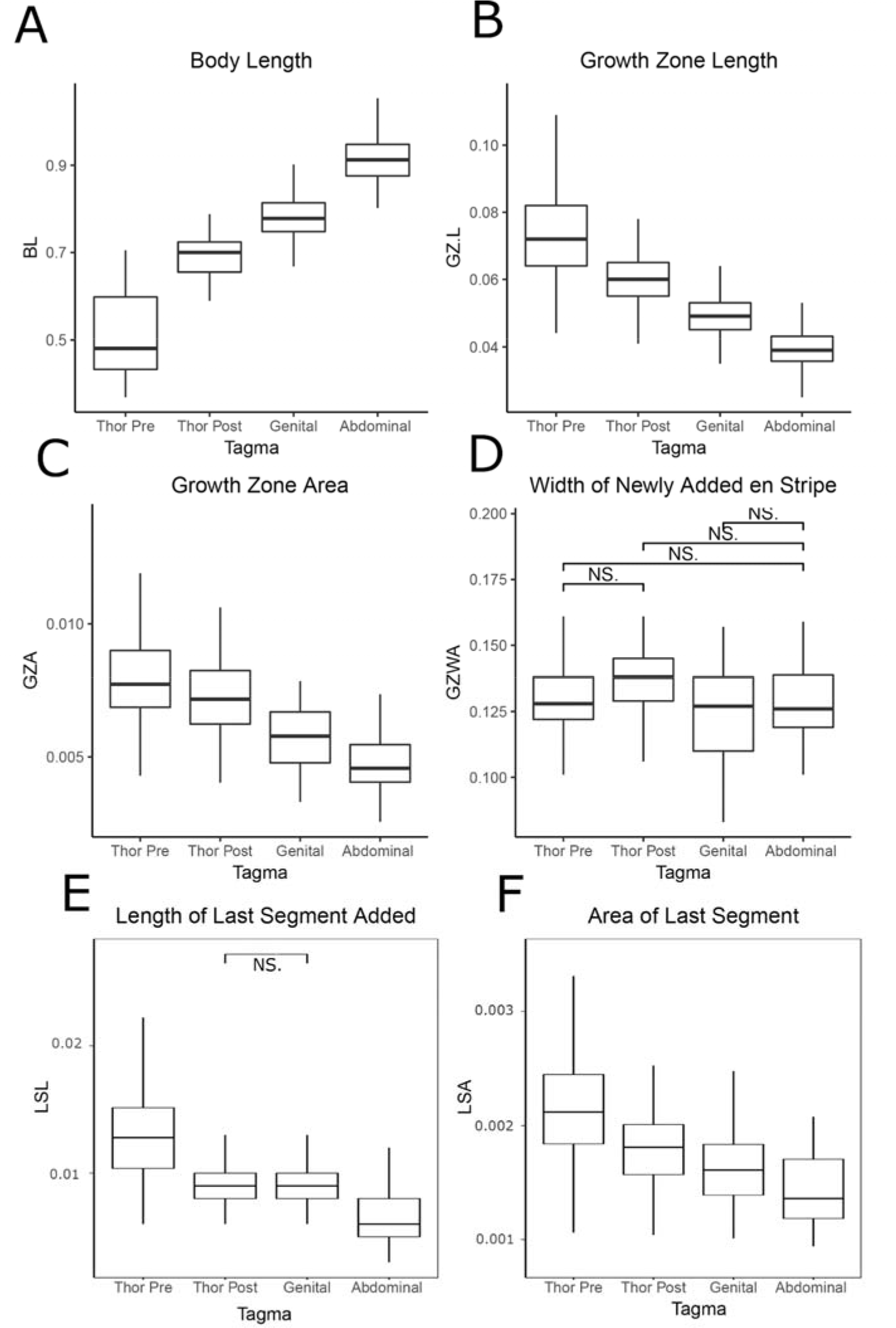
Tagma level differences in *Thamnocephalus* morphometric measurements. Tagma level differences (including pre- and post-molt thoracic ‘tagma’ identified from PCA; see Figure 5) are shown for body length (A), growth zone length (B) and area (C), the width of the newly added *en* stripe (D), last segment length (E) and area (F). All comparisons are significantly different (Tukey’s HSD; p<0.05) unless otherwise notated with “NS.”. The y-axes are measured in mm. Thor Pre= thoracic pre-molt; Thor Post= thoracic post-molt. Note- actual pdf is available <u>here</u>

**Supplementary Figure S4.**
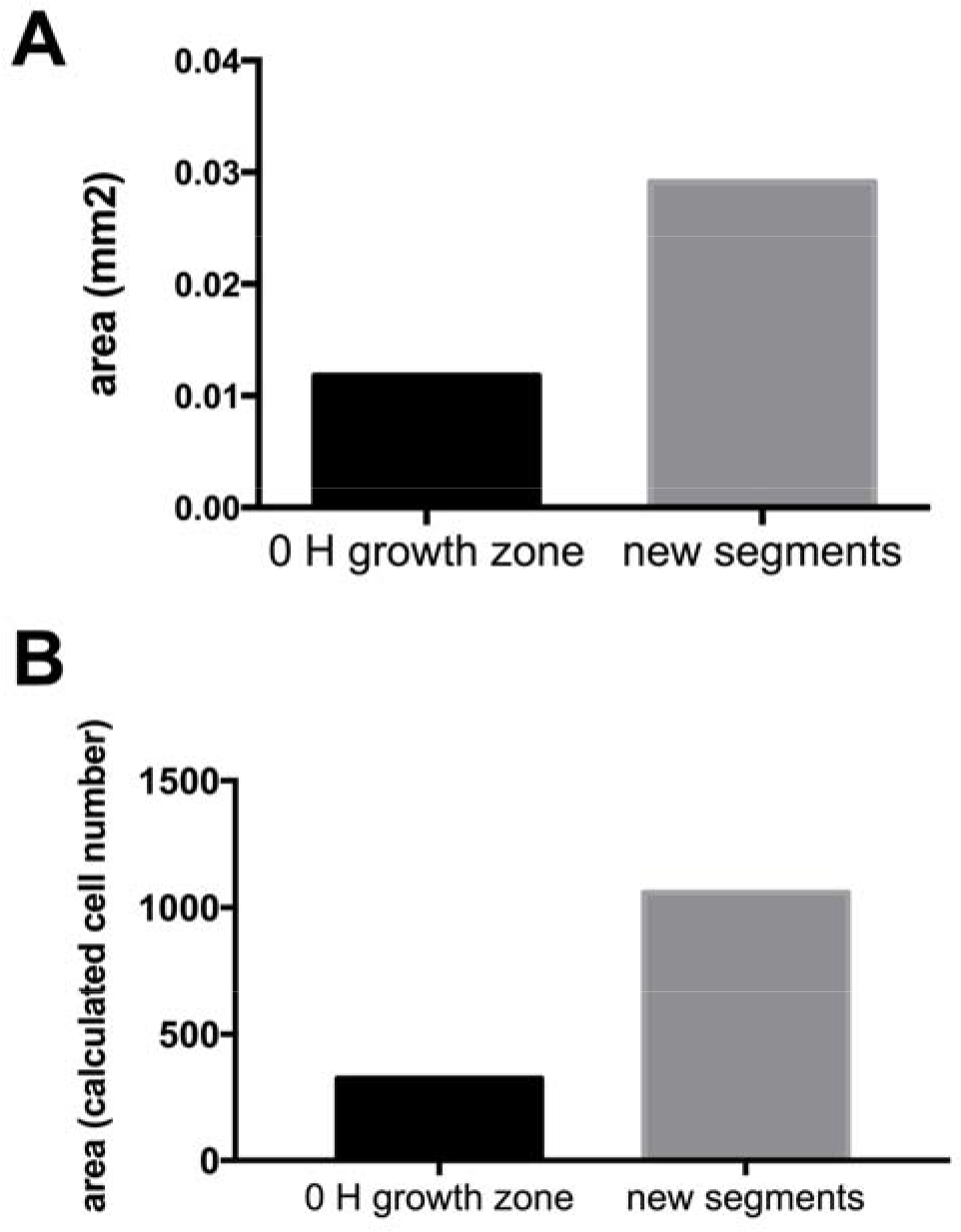
Comparison of tissue needed versus available in the growth zone to make new segments in a hatching larva. The average size of the initial growth zone upon hatching *versus* the area required to make all additional segments, where the latter is calculated based on the sum of each newly added segment over the measured course of development. A is based on linear measures, B is based on cell counts.

**Supplementary Figure S5.**
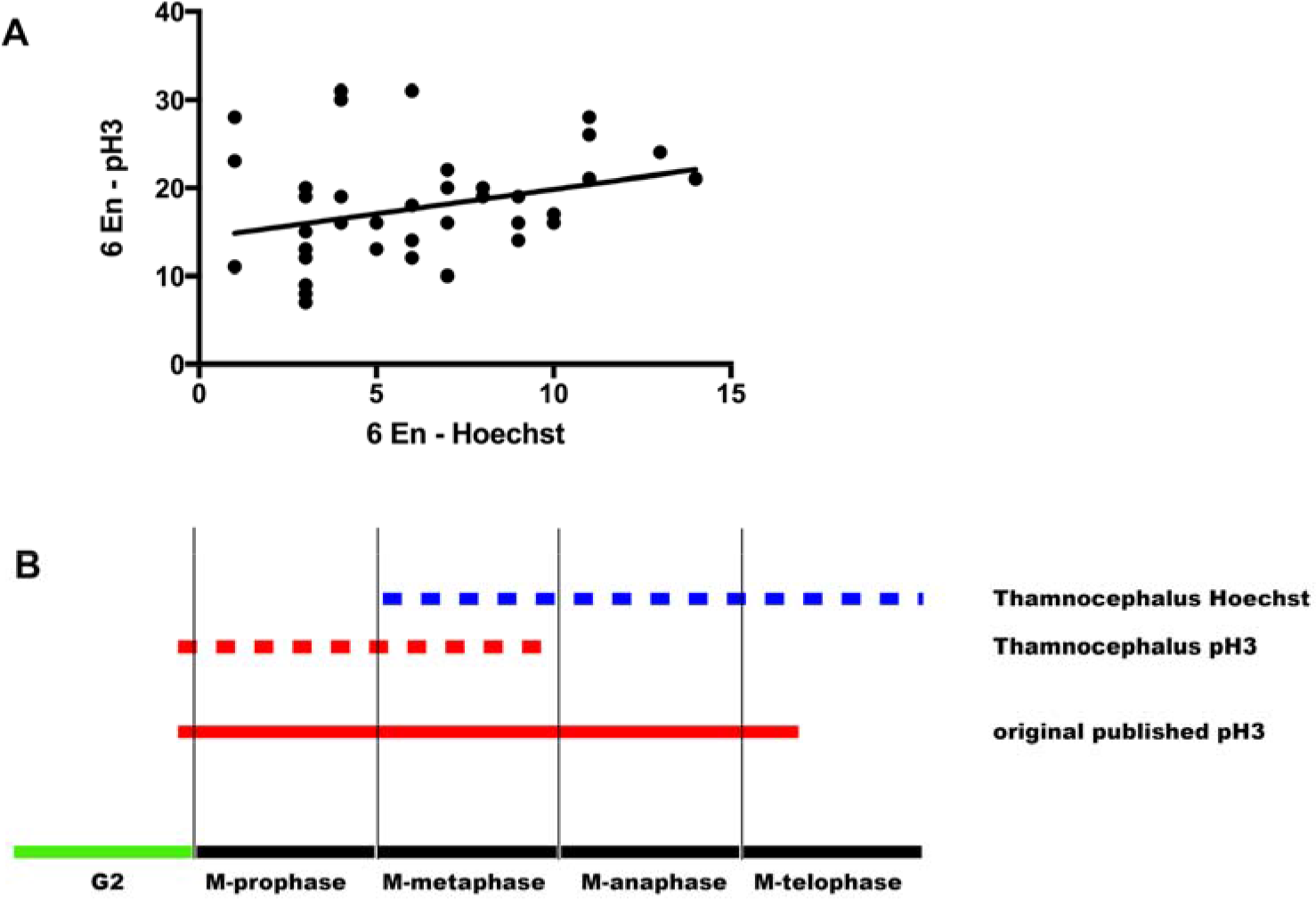
Correlation of pH3 and Hoechst mitosis counts and cell cycle expression. A. pH3 and Hoechst count correlation for 6 EN animals. We find low correlation at all developmental stages. B. Expression of growth zone pH3 and Hoechst in relation to cell cycle progression. Although pH3 is reported to be expressed throughout M-phase (red line), we find *Thamnocephalus* pH3 to be expressed early in M-phase (red dotted line). By comparison, mitosis counts using Hoechst only score cells in late M-phase.

**Supplementary Figure S6.**
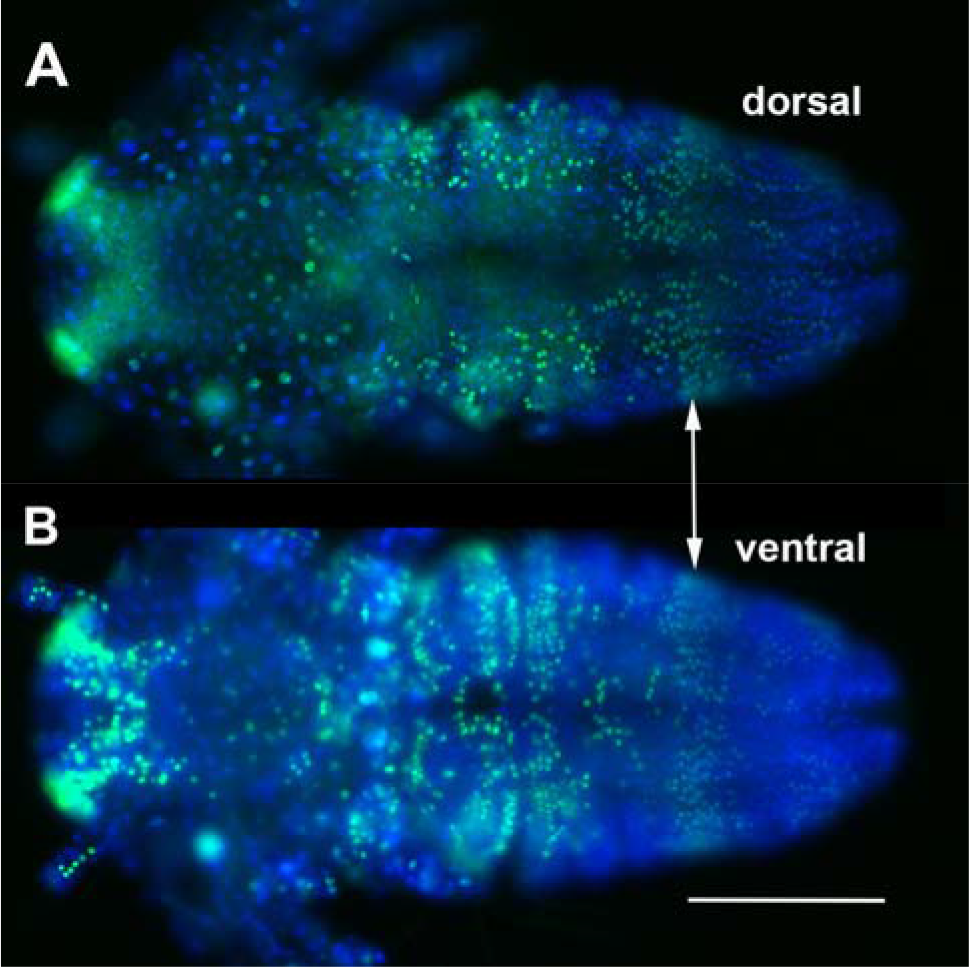
Comparison of dorsal and ventral cell dynamics in *Thamnocephalus* larvae, visualized by EdU incorporation. The pattern of Edu and all growth zone measures carry around to the dorsal side of the larvae (shown in focus in A). Focusing through the same specimen shows the normal pattern we describe in the text (B, cells out of focus due to being viewed through dorsal tissue). This corresponding patterning justifies restricting our measures and calculations to the ventral surface since we focus on changes in dimension and other relative features, not absolute measures.

**Supplementary Table 1.**
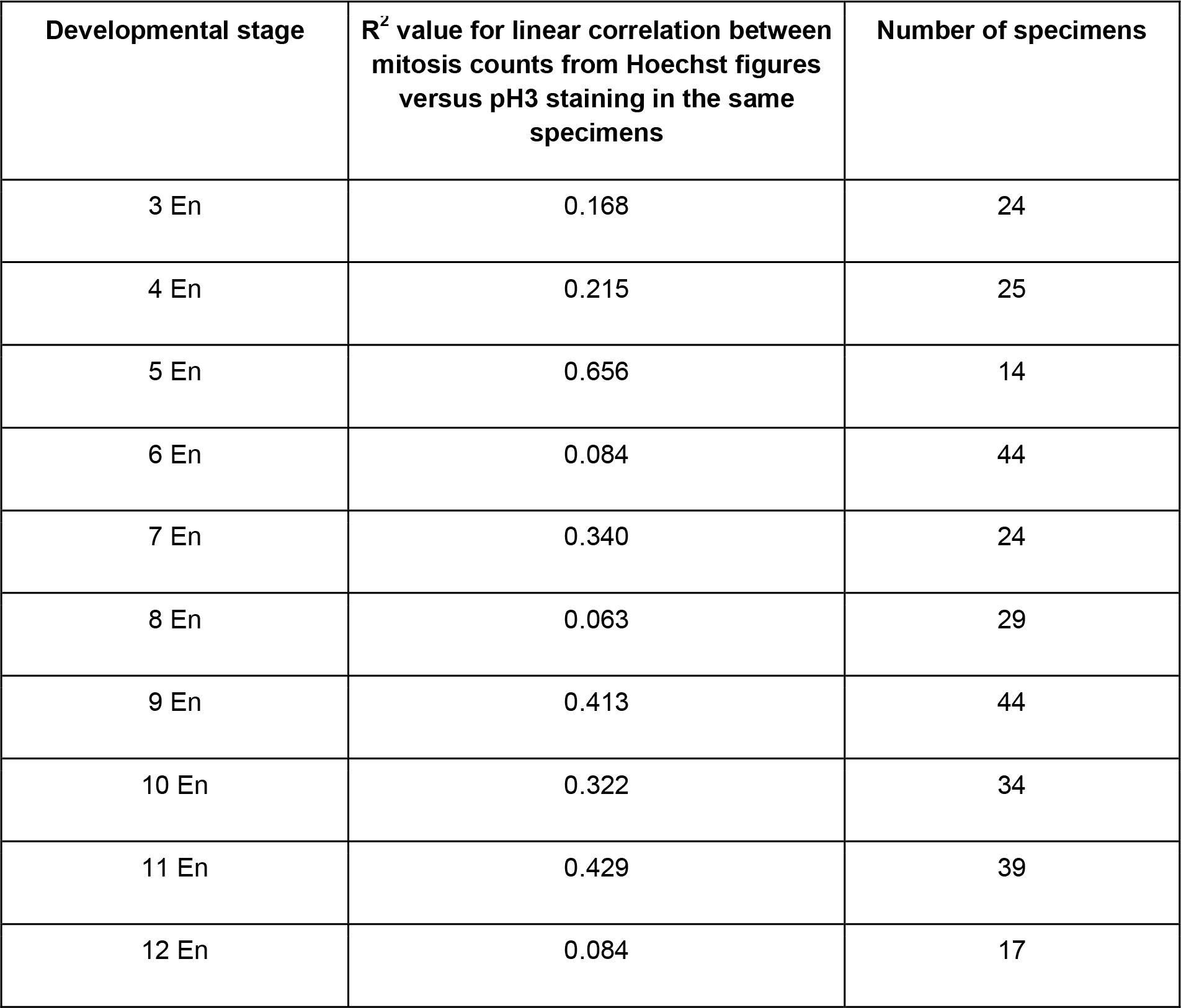
**Correlation between Hoechst and pH3 mitosis counts within the same individual.** For all developmental stages that have both Hoechst and pH3 data, the linear correlation and number of specimens is given.

| Developmental stage | R<sup>2</sup> value for linear correlation between mitosis counts from Hoechst figures versus pH3 staining in the same specimens | Number of specimens |
| --- | --- | --- |
| 3 En | 0.168 | 24 |
| 4 En | 0.215 | 25 |
| 5 En | 0.656 | 14 |
| 6 En | 0.084 | 44 |
| 7 En | 0.340 | 24 |
| 8 En | 0.063 | 29 |
| 9 En | 0.413 | 44 |
| 10 En | 0.322 | 34 |
| 11 En | 0.429 | 39 |
| 12 En | 0.084 | 17 |

## Author Contributions

Conceptualization: LMN, ADC, TAW

Methodology: SJC, TAW

Validation: SJC, TAW

Formal analysis: ND, SJC, TAW

Investigation: ND, SJC

Resources: TAW

Data curation: SJC, TAW

Writing - original draft: SJC, TAW

Writing - review & editing: ND, LMN, ADC, SJC, TAW

Visualization: ND, SJC, TAW

Supervision: TAW

Project administration: TAW

Funding acquisition: LMN, ADC, TAW

